# Flow cytometric isolation of drug-like conformational antibodies specific for amyloid fibrils

**DOI:** 10.1101/2023.07.04.547698

**Authors:** Alec A. Desai, Jennifer M. Zupancic, Hanna Trzeciakiewicz, Julia E. Gerson, Kelly N. DuBois, Mary E. Skinner, Lisa M. Sharkey, Nikki McArthur, Sean P. Ferris, Nemil N. Bhatt, Emily K. Makowski, Matthew D. Smith, Hongwei Chen, Jie Huang, Cynthia Jerez, Ravi S. Kane, Nicholas M. Kanaan, Henry L. Paulson, Peter M. Tessier

**Author notes:** To whom correspondence should be addressed: Peter M. Tessier Address: North Campus Research Complex, B10-179 2800 Plymouth Road University of Michigan Ann Arbor, MI 48109. co-first author (Alec A. Desai, Jennifer M. Zupancic).

## Abstract

Antibodies that recognize specific protein conformational states are broadly important for research, diagnostic and therapeutic applications, yet they are difficult to generate in a predictable and systematic manner using either immunization or *in vitro* antibody display methods. This problem is particularly severe for conformational antibodies that recognize insoluble antigens such as amyloid fibrils associated with many neurodegenerative disorders. Here we report a quantitative fluorescence-activated cell sorting (FACS) method for directly selecting high-quality conformational antibodies against different types of insoluble (amyloid fibril) antigens using a single, off-the-shelf human library. Our approach uses quantum dots functionalized with antibodies to capture insoluble antigens, and the resulting quantum dot conjugates are used in a similar manner as conventional soluble antigens for multi-parameter FACS selections. Notably, we find that this approach is robust for isolating high-quality conformational antibodies against tau and α-synuclein fibrils from the same human library with combinations of high affinity, high conformational specificity and, in some cases, low off-target binding that rival or exceed those of clinical-stage antibodies specific for tau (zagotenemab) and α-synuclein (cinpanemab). This approach is expected to enable conformational antibody selection and engineering against diverse types of protein aggregates and other insoluble antigens (e.g., membrane proteins) that are compatible with presentation on the surface of antibody-functionalized quantum dots.

## INTRODUCTION

Conformational antibodies are important for studying structure-function relationships for diverse types of proteins, including poorly soluble ones such as membrane^1, 2^ and amyloid-forming^3–5^ proteins. In the case of neurodegenerative diseases, conformational antibodies are critically important because of the large number of possible structures that can form for a single polypeptide assembled into diverse types of aggregates ranging from pre-fibrillar soluble oligomers to amyloid fibrils. Immunization^6–8^ and *in vitro* display technologies^9–11^, such as phage and yeast surface display, have been widely used to generate antibodies with conformational specificity against Aβ, tau, α-synuclein and other proteins linked to neurodegenerative diseases.

Despite this progress, the generation of high-quality conformational antibodies with strict conformational specificity for insoluble antigens such as protein aggregates remains largely a trial-and-error process, requires extensive experimentation and, in many cases, results in sub-optimal antibodies. In particular, most previously reported conformational antibodies against protein aggregates lack optimal combinations of three key properties, namely high affinity, strict conformational specificity, and low off-target binding^12, 13^. One key explanation for these sub-optimal combinations of binding properties is that it is challenging to select for all three antibody properties during the discovery process. This commonly results in antibodies generated against protein aggregates that i) bind linear peptide epitopes and lack conformational specificity, ii) display polyreactivity, and/or iii) possess modest affinity, especially for antibodies generated using *in vitro* display methods.

In principle, *in vitro* directed evolution methods such as yeast surface display should be able to address these challenges because of their ability to select for i) antibody binding at extremely low concentrations of aggregated antigen (high affinity), ii) a lack of antibody binding to non-aggregated antigen (high conformational specificity) and iii) a lack of binding to polyreactivity reagents (low antibody off-target binding). However, in practice, protein aggregates are typically insoluble antigens that are poorly compatible with quantitative cell sorting methods such as fluorescence-activated cell sorting (FACS). Instead, protein aggregates typically must be immobilized on surfaces such as magnetic beads for cell sorting via magnetic-activated cell sorting (MACS). The inability of MACS to perform multi-parameter, quantitative cell sorting leads to antibody selections that are strongly influenced both by the antibody expression level (e.g., display levels on the surface of yeast) and the intrinsic antibody affinity, which commonly leads to the enrichment of sub-optimal conformational antibodies and requires extensive secondary screening to isolate rare antibodies with strict conformational specificity, high affinity, and low off-target binding.

Here we report an unexpectedly simple approach for performing quantitative sorting of an off-the-shelf, yeast-displayed human antibody library against amyloid fibril aggregates (**Fig. 1**). Our approach is to conjugate capture antibodies (e.g., anti-tau antibodies) to quantum dots (QDs) and use the immunoconjugates to capture amyloid fibrils (e.g., tau fibrils). The resulting QD-amyloid conjugates are then used for quantitative, multi-parameter FACS sorting of a single, widely accessible human library to directly select antibodies with high affinity, strict conformational specificity and low polyreactivity. Herein, we demonstrate that this methodology can be used to generate conformational antibodies in a robust and predictable manner with combinations of binding properties that are similar to or better than those of clinical-stage antibodies.

**Figure 1.**
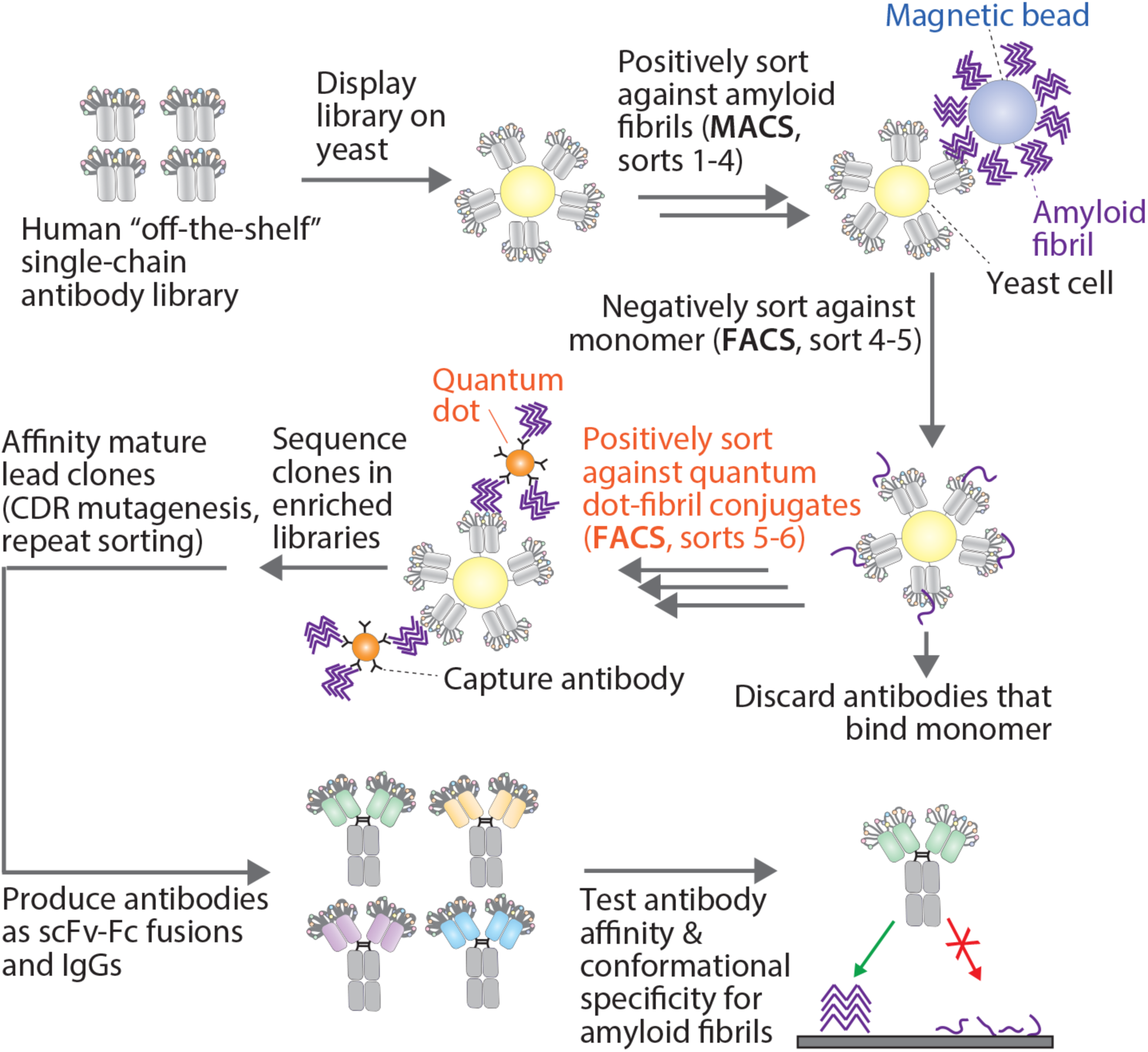
Overview of approach for selecting and affinity maturing anti-amyloid conformational antibodies. A previously reported human single-chain variable fragment (scFv) library^14^ was displayed on the surface of yeast and sorted positively for binding to amyloid fibrils (tau and α-synuclein) via magnetic-activated cell sorting (MACS) and negatively for a lack of binding to the monomeric form of the corresponding amyloid protein (tau or α-synuclein monomer) via fluorescence-activated cell sorting (FACS). The resulting enriched libraries were next positively sorted via FACS for binding to amyloid fibrils immobilized on quantum dot immunoconjugates. Quantum dot immunoconjugates were prepared using commercially-available quantum dots that enable the immobilization of a capture antibody (specific for tau or α-synuclein) on their surface. Next, the immunoconjugates were incubated with fibrils immediately prior to their use in antibody selections. Finally, the enriched antibody libraries were sequenced, and soluble antibody candidates were tested for their affinity and conformational specificity for amyloid aggregates using recombinant and biological samples. The schematic representations of magnetic beads, quantum dots, capture antibody, fibrils, monomer, yeast cells, and scFvs are not drawn to scale.

## RESULTS

### Generation of human conformational antibodies specific for tau amyloid fibrils

To evaluate the feasibility of quantitative antibody library sorting using QD-amyloid conjugates to generate human conformational antibodies in a simple and predictable manner, we first sorted an existing and widely used human antibody library against human tau fibrils. The single-chain (scFv) antibody library^14^ was sorted initially against the anti-Myc antibody to enrich the library for full-length antibodies. Next, the enriched library was sorted against tau aggregates by MACS for three rounds (**Fig. 2**; initial discovery, rounds 1-3). The percentage of collected cells increased from ∼0.04% (round 1) to ∼0.6% (round 3) over three consecutive rounds of MACS, indicating that the library was enriched for tau binding antibodies. Next, we performed a FACS negative selection (round 4) against disaggregated tau monomer to remove antibodies with low conformational specificity. Finally, a FACS positive selection (round 5) was performed against QD-tau aggregate conjugates to enrich for antibodies that bound to tau aggregates in a manner proportional to the antibody expression level. Notably, we detected a distinct population of antibody-displaying cells with affinity for tau fibrils that could easily be isolated using FACS.

**Figure 2.**
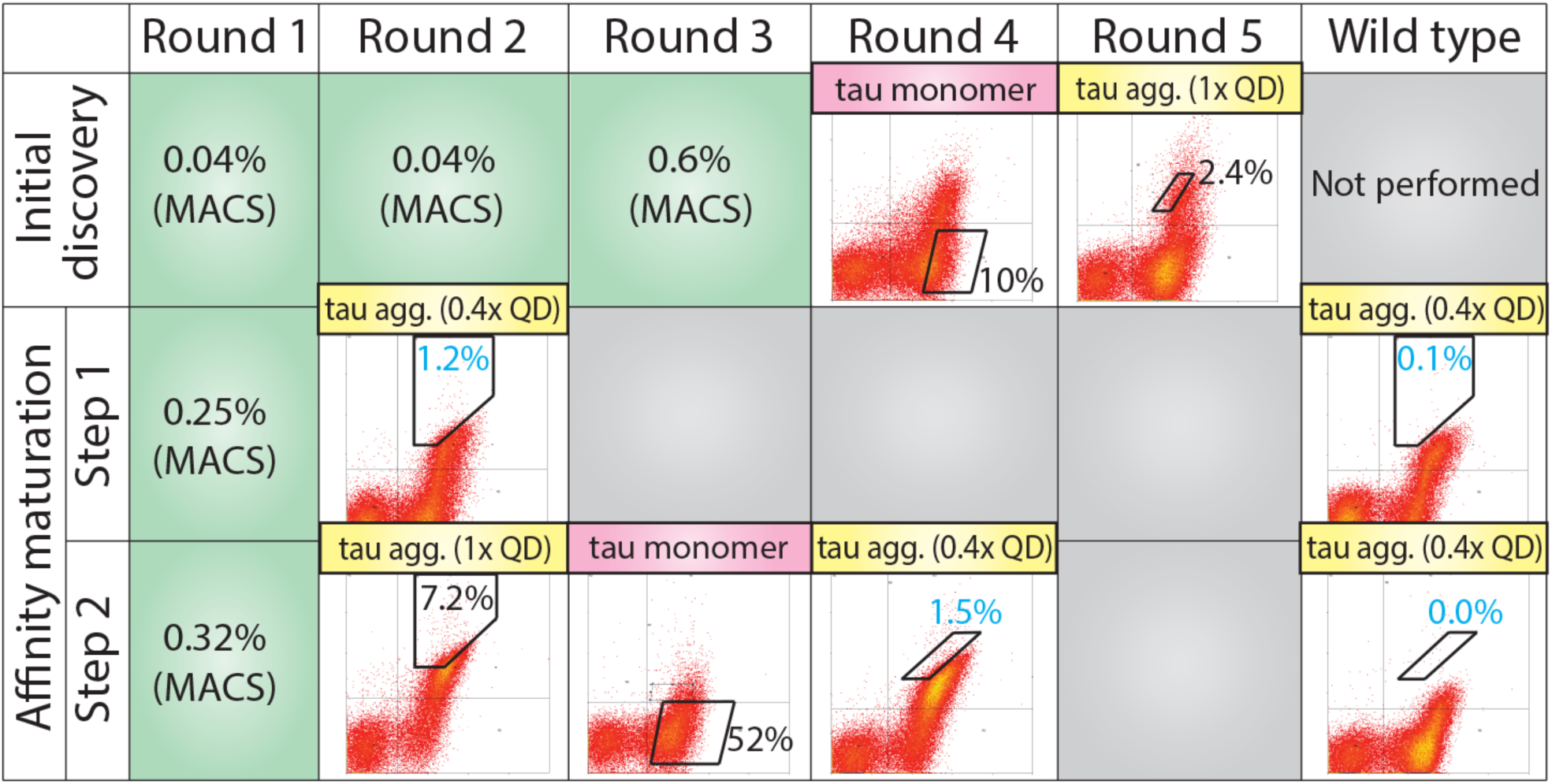
Summary of library sorting to enrich for tau conformational antibodies. In the initial discovery stage (top row), the scFv library was enriched three times against tau fibrils via MACS (round 1-3; percentage of cells collected is reported), depleted once against tau monomer to remove non-conformational antibodies via FACS (round 4), and enriched one more time against tau fibrils immobilized on quantum dot (QD) immunoconjugates via FACS (round 5). Next, a lead antibody clone was identified (ATA1) and mutagenized using soft randomization by mutating one CDR at a time for four CDRs, including heavy chain CDRs 1 and 2 and light chain CDRs 1 and 3 (four libraries). In step 1 of the affinity maturation (middle row), the libraries were enriched against tau fibrils using MACS (round 1) and QD conjugates using FACS (sort 2). Three of the mutated CDR libraries showed the strongest enrichment (HCDRs 1 and 2, LCDR3), including the HCDR2 library shown in the figure as a representative example (round 2 of step 1). In step 2 of the affinity maturation (bottom row), the three most promising CDR libraries were shuffled and enriched against tau fibrils using MACS (round 1) and QD conjugates using FACS (round 2), depleted once against tau monomer to remove non-conformational antibodies via FACS (round 3), and enriched one addition time against QD-tau fibril conjugates via FACS (round 4). The terms for tau in the figure refer to aggregated tau fibrils (tau agg.) and tau monomer. The percentages in the green boxes represent the % cells collected in the MACS sorts, while the percentages in the flow cytometry gates represent the % cells collected in the FACS sorts. The blue numbers represent the % of cells collected for the antibody libraries and wild-type samples that were evaluated at the same time, which can be directly compared.

We next Sanger sequenced the enriched library after round 5 and identified a lead tau antibody (ATA1; **Fig. S1**) for further characterization. ATA1 displayed specific binding to tau fibrils with intermediate affinity (EC_50_ of 6.3±1.5 nM), which was an order of magnitude weaker than the affinity of a clinical-stage tau conformational antibody (zagotenemab, EC_50_ of 0.5±0.10 nM; **Fig. 3A**). Encouragingly, ATA1 demonstrated high conformational specificity for tau fibrils, even when pre-incubated with a large excess of tau monomer (**Fig. 3B**). For example, at a 100-fold molar of excess of tau monomer relative to antibody, ATA1 retained 93% of its binding to tau fibrils. As controls, we confirmed that zagotenemab (conformational antibody) and Tau-5 (non-conformational antibody) retained most (83% for zagotenemab) or little (0.5% for Tau-5) of their binding to tau fibrils when pre-incubated with a 100-fold molar excess of tau monomer.

**Figure 3.**
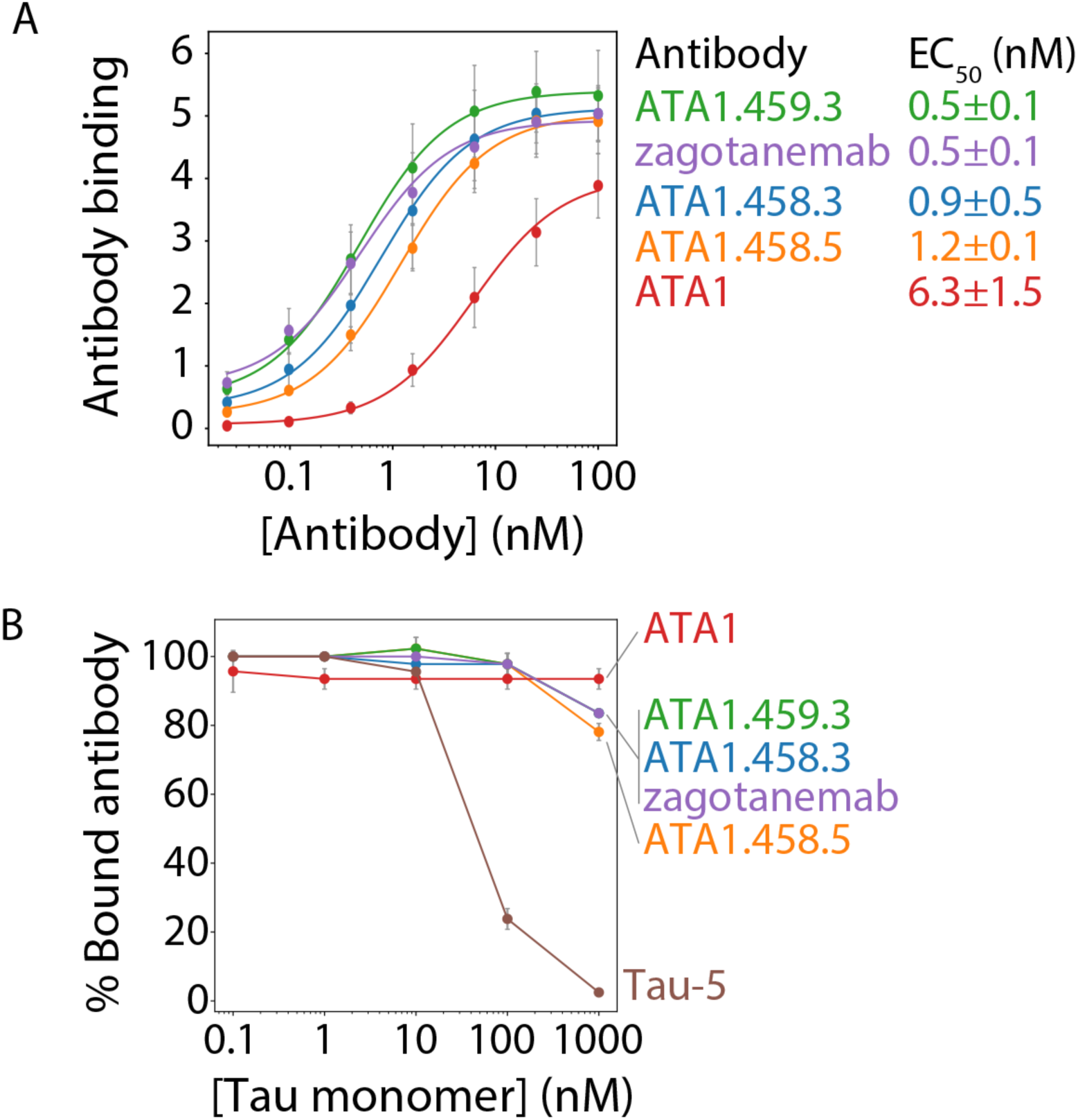
Affinity and conformational specificity of selected tau antibodies. (A) Tau antibodies were incubated with magnetic beads coated with tau fibrils, and the bound antibody signal was detected using fluorescently labeled anti-Fc secondary antibodies via flow cytometry. (B) Tau antibodies at a fixed concentration (10 nM) were pre-incubated with tau monomer (0-1000 nM), and then the mixtures were incubated with magnetic beads coated with tau fibrils. Bound antibody was subsequently detected using fluorescently labeled secondary antibody via flow cytometry, which is reported as a percentage of the antibody binding without tau monomer pre-incubation. In (A) and (B), the ATA1 series of antibodies are scFv-Fc fusion proteins, while zagotenemab and Tau-5 are IgG1s. The data are average of two independent experiments, and the errors are standard deviations.

These promising results for ATA1 motivated us to affinity mature it using a simple procedure that has been widely validated, namely soft randomization of single complementarity-determining regions (CDRs)^15, 16^. This procedure involves designing degenerate codons that sample the wild-type residue with a frequency of ∼50% and the other 19 amino acids with a cumulative frequency of ∼50%. We subjected four single CDRs to mutagenesis, namely HCDR1, HCDR2, LCDR1 and LCDR3. HCDR3 was not mutated because of its importance to antibody affinity, and LCDR2 was not mutated because of its relatively small size.

Each CDR library was separately sorted against tau aggregates in round 1 using MACS to reduce library diversity and enrich for binders (**Fig. 2**, affinity maturation step 1). Next, we performed FACS against QD-tau aggregate conjugates at a lower conjugate concentration (0.4x of what was used in round 5 of the initial discovery sorts) to enrich for higher affinity binders. This sorting resulted in three of four CDR libraries (HCDR1, HCDR2 and LCDR3) binding to QD-tau aggregate conjugates at higher levels than that for the parental antibody (ATA1). For example, we collected the enriched HCDR2 library in a gate with ∼1.2% of the antibody library-displaying cells that only includes 0.1% of the parental antibody-displaying cells.

Next, we sought to combine beneficial mutations from each of the three enriched CDR libraries (HCDR1, HCDR2 and LCDR3) to isolate antibodies with even higher affinities while maintaining high conformational specificity for tau aggregates (**Fig. 2**, affinity maturation step 2). To do this, we PCR amplified the diversified CDRs from each of the three enriched CDR libraries, shuffled them together using PCR, and constructed the full single-chain antibody genes. Next, the CDR-shuffled library was enriched for tau aggregate binders using MACS (round 1) and FACS (round 2). A negative FACS selection was performed against disaggregated tau monomer to remove non-conformational tau binders (round 3), and a final positive FACS selection was performed against QD-tau aggregate conjugates in round 4. The library was enriched to the point that we collected ∼1.5% of library-displaying cells in a gate that corresponded to 0.03% of the parental antibody-displaying cells.

We next evaluated antibody mutants from the enriched library by directly cloning them into Fc fusion plasmids for Sanger sequencing and soluble antibody expression. We produced 16 unique antibodies, purified them by Protein A chromatography, and characterized a subset of them via SDS-PAGE (**Fig. S2A**) and analytical size-exclusion chromatography (SEC; **Fig. S2B**). The single-chain antibodies showed high purity, including >90% monomer via SEC.

The affinity and conformational specificity of the affinity-matured antibodies were next evaluated in the same manner as for the parental and control antibodies (**Fig. 3**). Encouragingly, the affinity-matured clones showed increased affinities (EC_50_ values of 0.4-1 nM) relative to the parental ATA1 antibody (EC_50_ of 6.3±1.5 nM) and similar affinities as zagotenemab (EC_50_ of 0.5±0.1 nM; **Fig. 3A**). The conformational specificities of the affinity-matured antibodies were also similar to that for zagotenemab and slightly reduced relative to the parental ATA1 antibody (**Fig. 3B**). In particular, one of the affinity-matured variants (ATA1.459.3) displayed affinity and conformational specificity that was similar to that of zagotenemab.

We also evaluated the conformational specificity of the tau antibodies using a sandwich ELISA, which confirmed that both antibodies displayed high affinity for tau fibrils and little affinity for tau monomer (**Fig. S3**). These ELISA results were in stark contrast to those for a different N-terminal tau antibody (Tau-13), which bound strongly to both tau fibrils and monomer and lacked conformational specificity. We also used the same recombinant monomer and aggregated tau samples in denaturing western blotting assays, which revealed that ATA1.459.3 and zagotenemab similarly recognized denatured tau monomers and aggregates, suggesting that their reactivity with tau is conformation dependent (**Fig. S3**).

We next examined the epitope of ATA1.459.3 using both peptide scanning and mutational analysis of tau protein variants (**Figs. S4** and **S5**). We first performed a coarse scan of the ATA1.459.3 epitope using a peptide array that consisted of 15-amino acid tau peptides – which were offset from each other by 4 amino acids – and covered the entire full-length (HT40) tau sequence (**Fig. S4**). This initial scanning identified a region near the N-terminus of tau that was recognized by ATA1.459.3. We further refined the epitope with a more detailed peptide scan, which revealed that tau residues 16-TYGL-20 are necessary for binding of ATA1.459.3. Additionally, we confirmed these findings using tau protein fragments and mutants in an ELISA assay, which localized the epitope to tau residues 13-HAGTYGLGD-21 (**Fig. S5**), consistent with the minimal epitope of residues 16-TYGL-20.

We also evaluated the ability of the tau antibodies to recognize tau aggregates formed in the brains of tau transgenic (P301S) mice and human patients with Alzheimer’s disease (AD) or Progressive Supranuclear Palsy (PSP) (**Table S1**). First, immunodot blot and western blotting of homogenized brain samples from transgenic and wild-type (age matched control) mice revealed that ATA1.459.3 and zagotenemab bound primarily to transgenic samples (**Fig. S6**). Second, we performed immunofluorescence analysis on the mouse brain samples and observed that ATA1.459.3 specifically stained the tau transgenic brains, and this staining co-localized with phosphorylated tau (AT8) staining and weakly co-localized with total tau (Tau-5) staining (**Fig. S7**). Similar specificity of ATA1.459.3 was also confirmed using human brain sections (**Fig. 4**). The staining of ATA1.459.3 was specific for AD and PSP disease brain samples and co-localized with both phosphorylated and total tau immunostaining. These results collectively demonstrate that ATA1.459.3 selectively recognizes tau conformers formed *in vivo* for both transgenic animals and humans with disease pathology.

**Figure 4.**
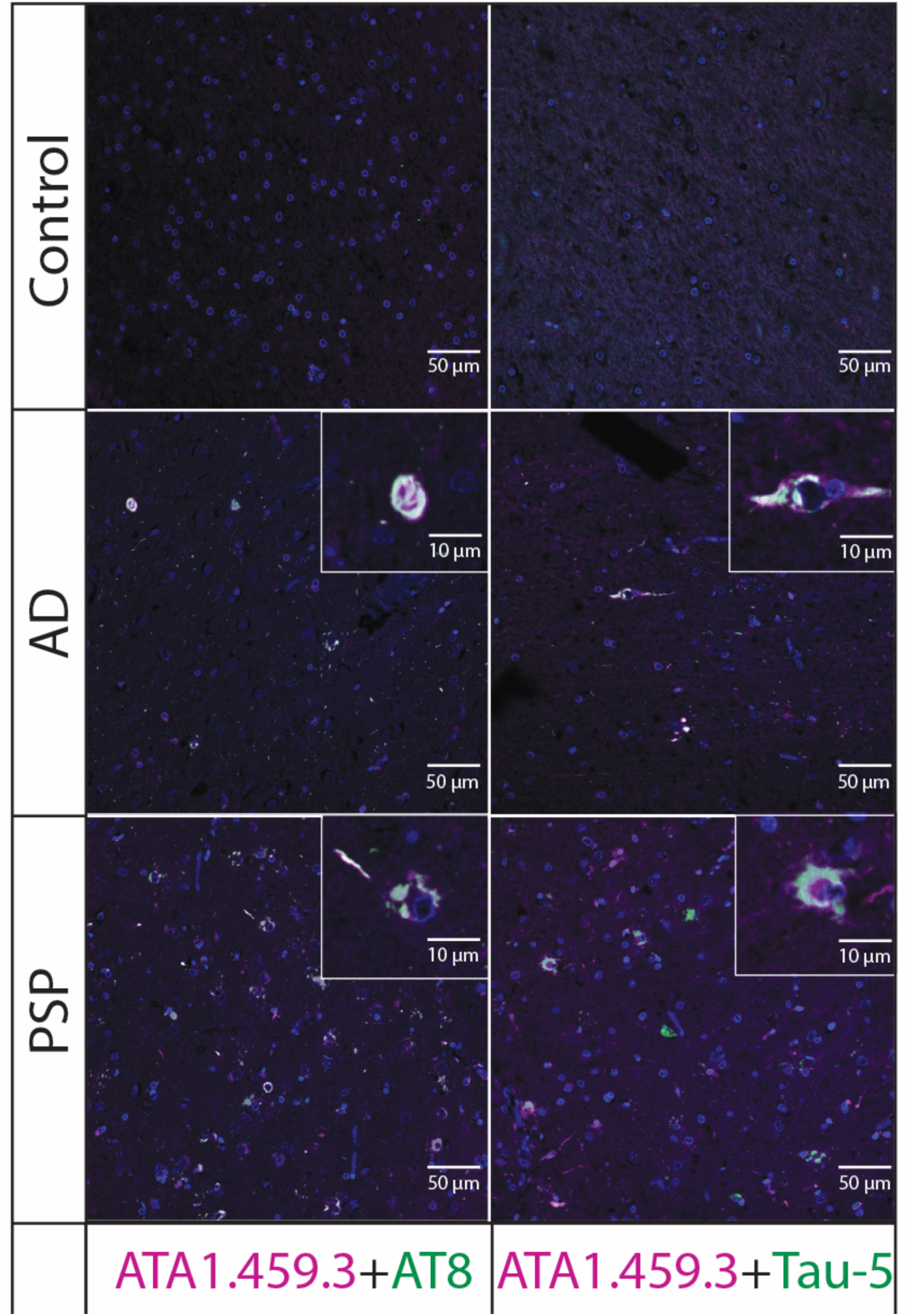
Tau immunofluorescence analysis of human brain tissues. Fixed human brain tissues from Alzheimer’s disease (AD) or Progressive Supranuclear Palsy (PSP) patients were co-stained with a conformational tau antibody (ATA1.459.3, purple), a phospho-tau (AT8, green) or pan-tau (Tau-5, green) antibody, and DAPI (blue). ATA1.459.3 was detected with anti-human Fc Alexa Fluor 647, and AT8 or Tau-5 were detected with anti-mouse Alexa Fluor 488. Merged staining results (with DAPI) are shown. The scale bars within the images are approximately 50 µm wide, and the scale bars within the insets are approximately 10 µm wide. ATA1.459.3 is an scFv-Fc fusion protein, while AT8 and Tau-5 are IgGs.

Finally, we also evaluated the ability of ATA1.459.3 to detect tau in human brain lysates using samples from Alzheimer’s disease and control tissues and prepared using both non-denaturing and denaturing conditions. We probed antibody binding using an assay format (sandwich ELISA) that does not require denaturation of the brain lysates. We observed much stronger binding of both ATA1.459.3 and zagotenemab to lysate from AD brain samples than that from control brain samples (**Fig. S8**). This contrasts with the results from the tau antibody that recognizes both monomers and aggregates (Tau-13), which displayed similar binding to lysates from AD and control brain samples. To further test the conformational dependence of antibody reactivity, we analyzed the binding of ATA1.459.3 and zagotenemab to the same brain lysates using denaturing conditions via western blotting, which revealed that both antibodies recognize multiple isoforms of tau in Alzheimer’s disease and control tissue (**Fig. S9**). Together, these results demonstrate that the antibodies are conformation dependent and detect an AD-related conformation of tau that is not present in control brains.

### Generation of human conformational antibodies specific for α-synuclein amyloid fibrils

We reasoned our discovery approach would be generally applicable to discovering human conformational antibodies specific for other types of amyloid fibrils. Therefore, we next evaluated the feasibility of using the same procedure, including using the same human antibody library, to select conformational antibodies specific for α-synuclein fibrils linked to Parkinson’s disease. The yeast-displayed scFv library was first enriched via MACS against immobilized α-synuclein fibrils (four rounds). During these sorts, the percentage of collected cells increased from 0.02% in round 1 to 0.4% in round 4 (**Fig. S10**, initial discovery). We next performed a negative selection against α-synuclein monomer via FACS (round 5) and a positive selection against QD-α-synuclein fibril conjugates via FACS (round 6). The output of round 6 was sequenced, which led to the isolation of a lead clone (aS2.1; **Fig. S11**). We produced this antibody in the IgG format and found that it possessed intermediate affinity (78.2±4.3 nM) relative to the clinical-stage antibody cinpanemab (9.5±2.6 nM; **Fig. 5A**). Encouragingly, aS2.1 displayed high conformational specificity for α-synuclein fibrils even when pre-incubated with 100-fold molar excess of α-synuclein monomer (∼90% of fibril binding activity), which was much higher than the corresponding conformational specificity of cinpanemab (∼44% of fibril binding activity; **Fig. 5B**).

**Figure 5.**
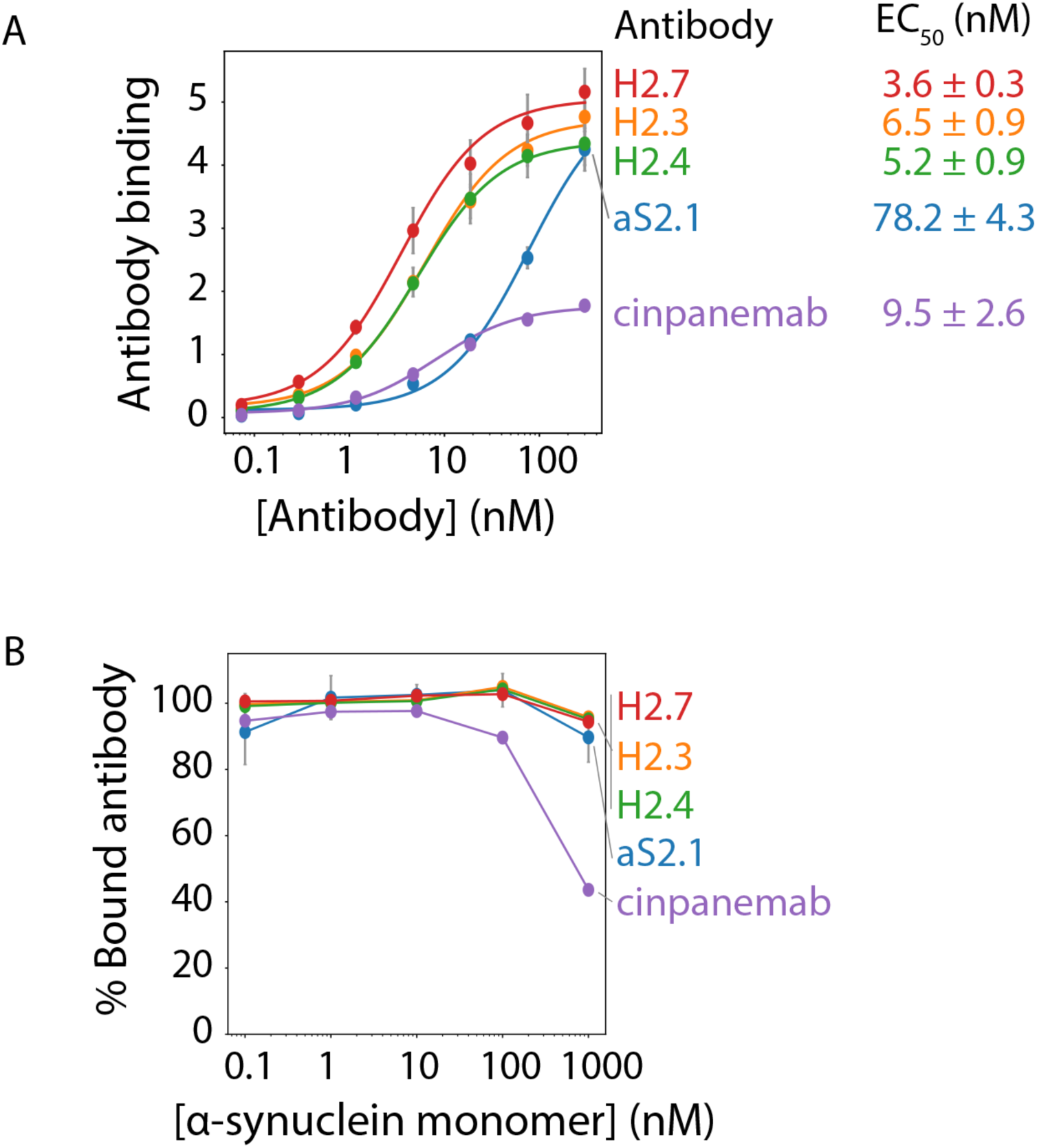
Affinity and conformational specificity analysis of α-synuclein antibodies. (A) α-synuclein antibodies (IgG1s) were incubated with Dynabeads coated with α-synuclein fibrils, and their binding was detected with fluorescently labeled anti-Fc secondary antibodies via flow cytometry. (B) α-synuclein antibodies at a fixed concentration (10 nM) were pre-incubated with α-synuclein monomer (0-1000 nM) followed by incubation with Dynabeads coated with α-synuclein aggregates. Antibodies bound to fibril-coated beads were detected by flow cytometry. The percentage of antibody binding to α-synuclein fibrils was calculated as the binding signal in the presence of α-synuclein monomer pre-incubation (0.1-1000 nM) relative to the control without pre-incubation. All antibodies were tested in IgG format with a human IgG1 Fc. The data are averages of two independent repeats, and the error bars are standard deviations.

We next affinity matured aS2.1 using a similar procedure as for the lead antibody isolated for tau (ATA1), as summarized in the Methods section and **Fig. S10**. We initially prepared soft mutagenesis libraries focused on each of the six individual CDR sequences for aS2.1. Results from the initial library sorting demonstrated that the HCDR2 library resulted in the greatest apparent improvement in affinity, and we thus chose to focus our analysis on clones selected from this library. This led to the isolation of three antibody variants (H2.3, H2.4 and H2.7) with mutations only in HCDR2 (**Fig. S11**). The IgGs were expressed in HEK293-6E cells, purified via Protein A chromatography, and their purity was examined by SDS-PAGE and SEC (**Fig. S12**). All antibodies showed high purity, including >90% monomer by SEC.

Notably, the three α-synuclein antibody mutants displayed both higher affinity (**Fig. 5A**) and conformational specificity (**Fig. 5B**) than cinpanemab using a flow cytometry assay. We also observed similar results for one of our selected mutants (H2.7) in comparison to cinpanemab using a non-denaturing, sandwich ELISA (**Fig. S13**). Importantly, H2.7 recognized both denatured monomers and aggregates, as detected via western blotting, revealing that its reactivity with α-synuclein is conformation dependent.

Immunoblot analysis revealed the matured antibodies also selectively recognized brain lysates from α-synuclein transgenic (A53T) mice relative to those for wild-type mice, especially for H2.7, in a manner similar to that for cinpanemab (**Fig. S14**). We also confirmed that H2.7 selectively immunostains human brain samples associated with Parkinson’s and Diffuse Lewy Body diseases relative to non-diseased brains (**Fig. 6**). Immunofluorescence analysis revealed co-staining of H2.7 with antibodies specific for phosphorylated α-synuclein (pSyn129) and total α-synuclein (LB509). We also examined H2.7 and cinpanemab binding to human brain samples via western blotting (**Fig. S15**) and found that both antibodies recognize α-synuclein in Lewy body dementia and control lysates.

**Figure 6.**
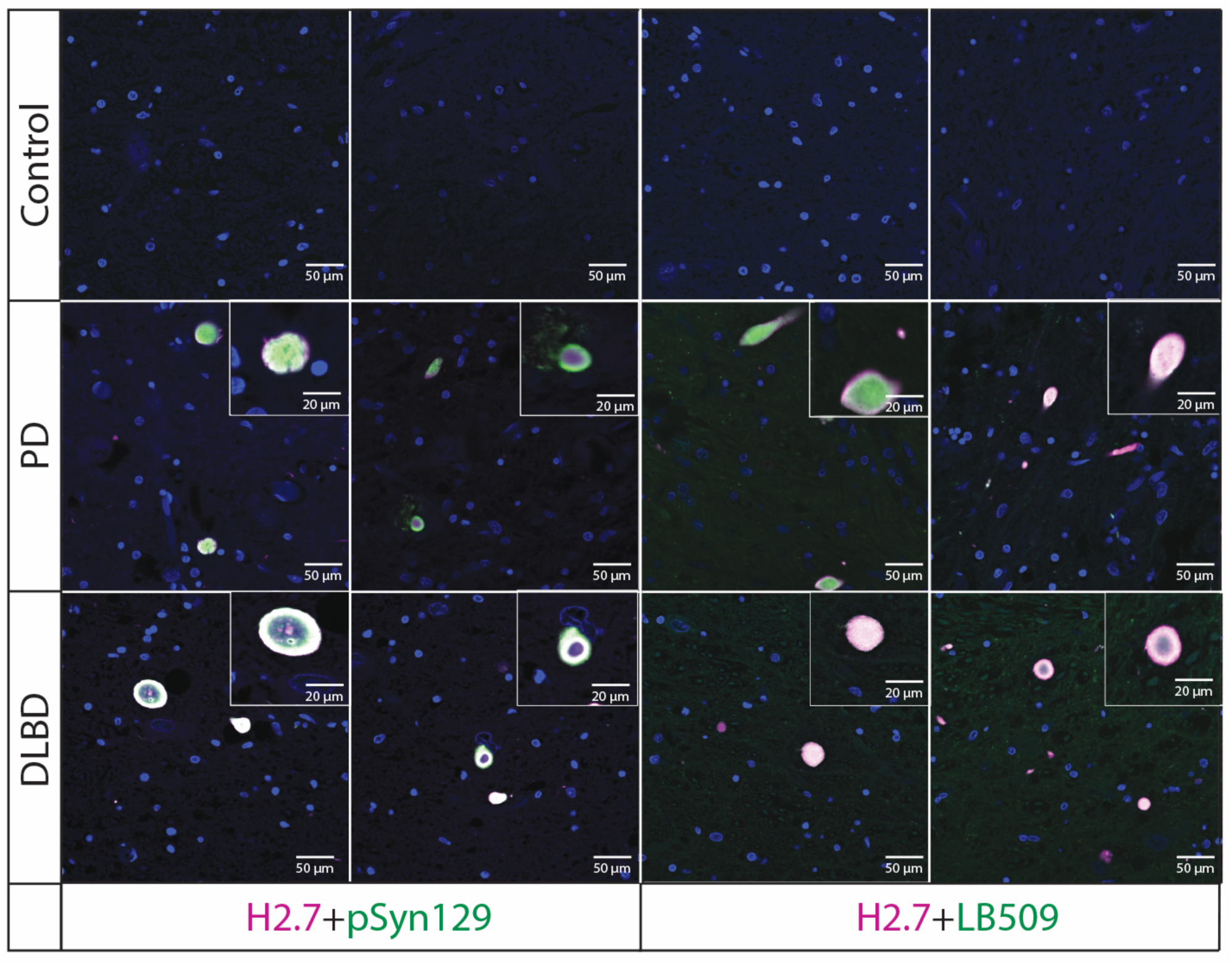
α-synuclein immunofluorescence analysis of human brain tissues. Fixed human brain tissues from Parkinson’s disease (PD) or Diffuse Lewy body disease (DLBD) patients were co-stained with H2.7 (conformational α-synuclein antibody) and EP1536Y (phospho-antibody specific for pSyn129) or LB509 (pan-synuclein) antibody. H2.7 was detected with anti-human Fc Alexa Fluor 647 (purple), LB509 was detected with anti-mouse Alexa Fluor 488 (green), and pSyn129 was detected with anti-rabbit Alexa Fluor 488 (green). Merged staining results with DAPI (blue) are shown. The scale bars within the images are approximately 50 μm wide, and the scale bars within the insets are approximately 20 μm wide. All antibodies were tested in IgG format.

### Tau and α-synuclein conformational antibodies selectively recognize their cognate aggregates

We next tested if the antibodies generated in this study display strict sequence specificity, as some previously reported conformational antibodies recognize a common conformational motif in aggregates formed from different proteins^8, 17^ while others have been reported which are both sequence- and conformation-specific^5, 18^. We first performed immunohistochemistry to analyze the binding of our antibodies to aggregates in human AD brain tissue that contains both tau and α-synuclein aggregates. To perform this staining, consecutive brain sections were obtained to enable a direct comparison of different antibodies binding to similar aggregates in each brain sample. Notably, we observe that our tau antibody (ATA1.459.3) strongly stains similar tau aggregates as those recognized by zagotenemab, including dystrophic neurites and neutrophil threads (**Fig. 7**). While multiple observed locations within the stained tissue contained both tau and α-synculein aggregates, ATA1.459.3 and zagotenemab did not recognize α-synculein aggregates (Lewy bodies) in these same regions. Our α-synuclein (H2.7) antibody strongly stained Lewy bodies, and this staining was similar to that observed for cinpanemab. However, neither α-synuclein antibody showed staining of nearby tau aggregates. Further, both conformational tau antibodies showed different staining patterns compared with a total tau antibody that recognizes both tau monomer and aggregates (Tau-5), and both conformational α-synuclein antibodies showed different staining patterns than a total α-synuclein antibody that recognizes both α-synuclein monomer and aggregates (LB509). Finally, the immunohistochemistry results for ATA1.459.3 (in AD tissue) and H2.7 (in PD tissue) were also evaluated using fixed free-floating sections that do not require antigen retrieval, which also confirmed specific antibody staining of disease-associated conformers (**Fig. S16**) while no antibody staining was observed in control tissue.

**Figure 7.**
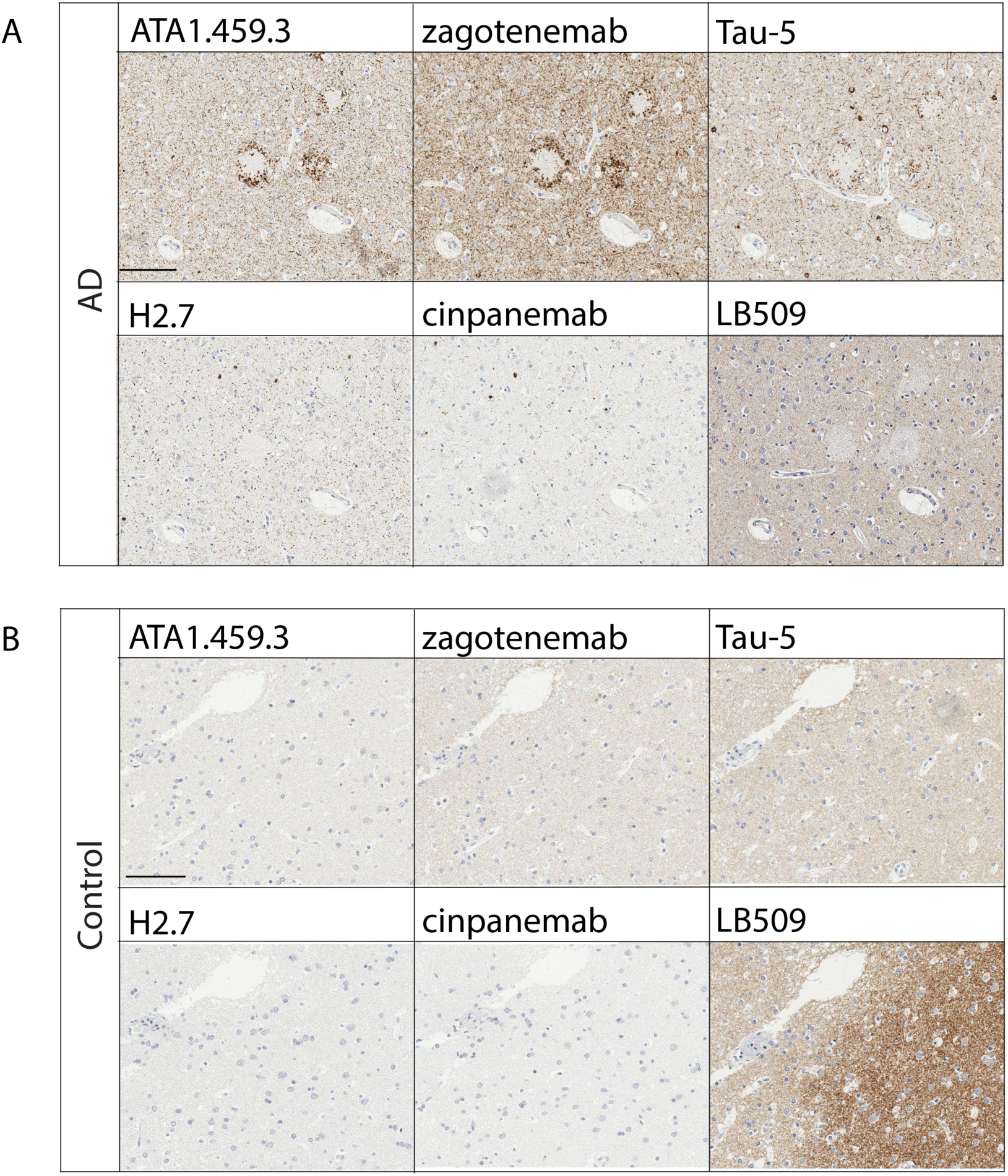
Immunohistochemical staining of human brain tissues. Fixed brain tissue from (A) Alzheimer’s disease (AD) samples in which both tau and α-synuclein aggregates were present and (B) control samples in which aggregates of both proteins were largely absent was stained with conformational and control antibodies. (A) Staining of AD tissue with ATA1.459.3 and zagotenemab shows similar staining of tau aggregates (distrophic neutrites and neutrophil threads) by both antibodies. Staining of the same tissue with H2.7 and cinpanemab shows similar staining of α-synuclein aggregates (Lewy bodies) by both antibodies. Images display the same region of consecutive sections from the frontal cortex from the same tissue in which both tau and α-synuclein aggregates are present. However, staining of α-synuclein aggregates by either of the conformational tau antibodies or staining of tau aggregates by either of the α-synuclein conformational antibodies was not observed. Control tau (Tau-5) and α-synuclein (LB509) antibodies were also included for representative staining of these regions by non-conformational antibodies. (B) Both ATA1.459.3 and H2.7 showed no observable staining when tested using control tissue. ATA1.459.3 was examined as an scFv-Fc fusion with a human IgG1 Fc. H2.7, cinpanemab, and zagotenemab were analyzed in human IgG1 format. Tau-5 and AT8 are IgGs with mouse Fc.

Additionally, we performed immunoblotting to further evaluate the binding of our antibodies to multiple forms of tau and α-synuclein (**Fig. S17**). This analysis, which involves direct immobilization of tau and α-synuclein conformers on immunoblots and can lead to denaturation and loss of conformational epitopes, is meant to evaluate sequence specificity of the antibodies. For ATA1.459.3, we observe antibody recognition of fibrils of human tau isoforms in which the N-terminus is intact (2N4R and 0N3R isoforms), while no binding was observed to mouse tau fibrils. The latter finding agrees with our identification of an N-terminal tau epitope because the tau sequences differ between human (residues 16-TYGL-20) and mouse (16-<u>D</u>Y<u>T</u>L-20) tau in this region. We also observe that zagotenemab recognizes similar tau conformers as ATA1.459.3. Similarly, we observe that H2.7 and cinpanemab recognize multiple α-synuclein conformers (fibrils and oligomers; **Fig. S18**), including human and mouse fibrils (**Fig. S17**). Finally, we observe minimal cross-reactivity of the tau antibodies (ATA1.459.3 or zagotenemab) with α-synuclein conformers, and no observable cross-reactivity of the α-synuclein antibodies (H2.7 or cinpanemab) for any of the tau conformers.

### Biophysical characterization of human conformational antibodies

To be used as therapeutic or diagnostic agents, conformational antibodies also need to possess a combination of favorable biophysical properties, including i) high stability, ii) high specificity (i.e., low off-target binding), and iii) low self-association. Therefore, we evaluated the biophysical properties of our antibodies using a panel of assays. First, we examined the specificity of the tau and α-synuclein antibodies by assaying their interactions with a polyspecificity reagent, namely soluble membrane proteins from CHO cells^19^, in comparison to clinical-stage antibodies which have been shown to have high (emibetuzumab) and low (elotuzumab) non-specific binding^19, 20^. The lead tau antibody (ATA1) as well as its affinity matured variants (ATA1.458.3, ATA1.458.5, and ATA1.459.3) showed low non-specific binding to soluble membrane proteins (**Fig. 8A**). In contrast, Tau-5 showed moderate non-specific binding and zagotenemab displayed high non-specific binding. For the α-synuclein antibodies, the lead antibody (aS2.1) and its affinity matured variants (H2.3, H2.4 and H2.7) demonstrated moderate to high non-specific binding, which was modestly higher than the level of non-specific binding for cinpanemab.

**Figure 8.**
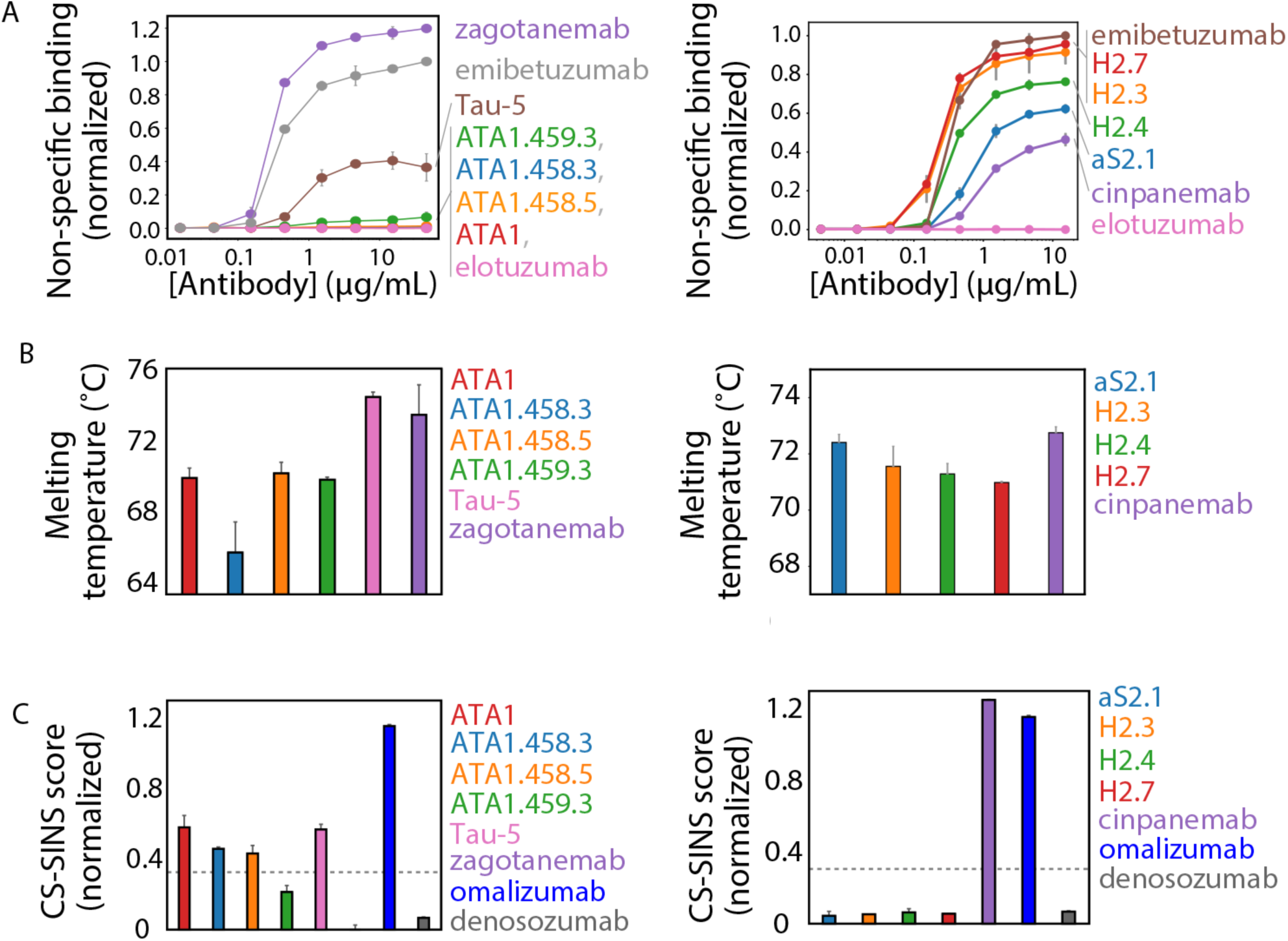
Biophysical characterization of tau and α-synuclein antibodies. (A) Non-specific binding of tau and α-synuclein antibodies to biotinylated soluble membrane proteins from CHO cells was analyzed by flow cytometry. The antibodies were immobilized on Protein A-Dynabeads, and binding of the soluble membrane proteins was detected via streptavidin Alexa Fluor 647. The binding signals were normalized between the signals for emibetuzumab (high non-specific binding) and elotuzumab (low non-specific binding). (B) Antibody melting temperatures (first unfolding transition) were analyzed by differential scanning fluorimetry. (C) Self-association of antibodies was analyzed by charge-stabilized self-interaction nanoparticle spectroscopy (CS-SINS). The dashed line represents a CS-SINS score of 0.35, and values <0.35 indicate low levels of self-association^23^. Tau and α-synuclein antibodies were analyzed in comparison to omalizumab (high self-association) and denosozmab (low self-association), and all data is normalized to a previously published scale. In (A-C), the data are averages of 2-3 independent repeats, and the error bars are standard deviations. In (A-C), the ATA1 series of antibodies are scFv-Fc fusion proteins, and all other antibodies are IgG1s.

Next, we analyzed the folding stability of the tau and α-synuclein antibodies to evaluate whether the lead or affinity-matured antibodies possess sub-optimal stabilities, such as melting temperatures (*T_m_*) below 65 °C (**Fig. 8B**). Encouragingly, the lead tau antibody (ATA1) displayed a relatively high melting temperature (*T_m_* of 69.8±0.6 °C), and this was similar for the affinity-matured variants (66-71 °C). The two IgGs (Tau-5 and zagotenemab) showed modestly higher stability (*T_m_* values of 73-74 °C), as expected for IgGs due to their stabilizing constant (C_H_1 and C_L_) regions that are absent in the scFv-Fc fusion proteins for ATA1 and the affinity-matured variants. We observed similarly encouraging results for the α-synuclein antibodies, as the lead antibody (aS2.1, formatted as an IgG) displayed high stability (*T_m_* of 72.2± 0.1 °C), and the affinity-matured variants (*T_m_* values of 71.7-72.4 °C) and cinpanemab (*T_m_* of 74.3±0.7 °C) displayed similar stabilities.

Finally, we examined the level of self-association for the tau and α-synuclein antibodies in a common formulation condition used for therapeutic antibodies, namely pH 6 and 10 mM histidine (**Fig. 8C**)^21^. High levels of antibody self-association are linked to high antibody viscosity and opalescence in concentrated formulations, such as those required for therapeutic formulations^22–24^. Therefore, we analyzed this property using charge-stabilized self-interaction nanoparticle spectroscopy (CS-SINS), which identifies antibodies with low self-association and low risk for high viscosity and opalescence in concentrated formulations as those with CS-SINS scores <0.35^25^. Encouragingly, our best affinity-matured tau antibody (ATA1.459.3) and three affinity matured α-synuclein antibodies (H2.3, H2.4 and H2.7) display CS-SINS scores <0.35. In contrast, the clinical-stage antibodies display mixed results, as zagotenemab displays low self-association and cinpanemab displays a high self-association. Overall, these results reveal that our tau and α-synuclein antibodies possess a combination of biophysical properties that are similar or superior to those for clinical-stage antibodies, which demonstrates the high-quality nature of the antibodies generated in our work against different amyloid targets from a single human library without the need for immunization.

## DISCUSSION

We have demonstrated a quantitative method for generating and affinity maturing conformational antibodies against two different amyloid-forming proteins using the same off-the-shelf human library. While the majority of conformational antibodies have previously been isolated using *in vivo* discovery methods, such approaches provide limited control over the selection process and do not reliably produce the desired conformational specificity. Our approach builds on previous work using nonimmune *in vitro* libraries to perform library panning^10, 11, 26–35^ and MACS^13, 36, 37^ to isolate conformational antibodies. Both techniques provide some control over antigen presentation and concentration during sorting. The majority of previous studies using these methods employed either alternating rounds of positive selections against immobilized aggregates and negative selections against immobilized monomers^26, 36^ or selections against immobilized aggregates in the presence of a competitive form of the amyloid protein (e.g., monomers) in solution^11, 27^. Despite some success, these approaches yield variable and hard-to-predict results. We find that multiparameter, quantitative selections for both antibody expression and either QD-amyloid binding or a lack of monomer (e.g., tau or α-synuclein) binding, which cannot be simultaneously performed using library panning or MACS, are critical for reliably identifying conformational antibodies with both high affinity and specificity.

We have demonstrated the strength of this FACS-based technique by isolating high-affinity conformational antibodies directed toward both tau and α-synuclein fibrils. The properties of the antibodies isolated using this method should be considered in the context of previously reported anti-amyloid antibodies. For this purpose, we have directly compared the properties of our antibodies to those of clinical-stage antibodies directed toward the same targets. The affinities and conformational specificities of our best tau antibodies as scFv-Fc fusion proteins (ATA1.459.3, ATA1.458.3, and ATA1.459.5) are comparable to those of zagotenemab (**Figs. 3** and **S3**), a conformational tau antibody evaluated in clinical trials^38^. Interestingly, we find that the epitope recognized by ATA1.459.3 comprises residues 16-20 of tau (**Figs. S4** and **S5**), which differs from that of the reported epitope for zagotenemab (residues 7-9 and 313-322).

It is also notable that zagotenemab shows similar specificity as ATA1.459.3 in terms of immunodot blotting of transgenic mouse brain samples, although zagotenemab also shows high background at long exposure times, while our antibody (ATA.459.3) does not (**Fig. S6**). This observation agrees with our finding that zagotenemab shows a high level of non-specific binding to a polyspecificity reagent (soluble membrane proteins; **Fig. 8A**), while our tau antibodies show minimal non-specific binding. Like zagotenemab (which is an IgG), our scFv-Fc antibodies were also stably folded (melting temperatures >65 °C; **Fig. 8B**) and our best tau antibody (ATA1.459.3) shows low self-association (**Fig. 8C**). These favorable biophysical properties relative to zagotenemab, especially for the ATA1.459.3 antibody, are particularly exciting given the ability of this antibody to recognize pathological tau aggregates in brain samples from transgenic mice (**Fig. S4**) and humans with Alzheimer’s disease (**Figs. 4, 7 and S16**) and progressive supranuclear palsy (**Fig. 4**). It is also interesting that we observe binding of this antibody to both control and AD brain tissue lysate when the lysate is denatured (**Fig. S9**), while we only observe antibody binding to the AD samples when the lysate is not denatured (**Fig. S8**). This difference in antibody binding between denaturing and non-denaturing conditions has been observed previously for other tau conformational antibodies that recognize related but unique N-terminal tau epitopes in tau aggregates^4^.

Similarly, the properties of our α-synuclein antibodies should be compared to those for cinpanemab, a conformational α-synuclein antibody that was evaluated in clinical trials^39^. Three of our best α-synuclein antibodies (H2.3, H2.4 and H2.7) displayed similar or higher affinity and conformational specificity than cinpanemab (**Fig. 5**). Our antibodies were also stably folded like cinpanemab (*T_m_* values >70 °C; **Fig. 8B**), and they show much lower levels of self-association (**Fig. 8C**) and moderately higher levels of non-specific binding (**Fig. 8A**) than cinpanemab. These generally favorable biophysical properties of our antibodies relative to cinpanemab are also encouraging because our best antibody variants, such as H2.7, display highly specific binding to pathological α-synuclein species in human brain samples from both Parkinson’s and diffuse Lewy body disease samples (**Figs. 6** and **S16**) as well as AD tissue which contains Lewy bodies (**Fig. 7**).

## CONCLUSIONS

The method we have developed presents a number of exciting opportunities for future development. First, we used a single capture antibody conjugated to the QDs for each antigen. However, this conjugation procedure is readily extendable to alternate capture antibodies, including mixtures of antibodies or polyclonal antibodies. For example, polyclonal antibodies have been developed that are conformationally specific for amyloid oligomers^5, 8, 40^ and fibrils^17^. We speculate that such polyclonal antibodies could be conjugated to QDs and used to isolate monoclonal antibodies specific for various types of amyloidogenic conformers. Further, our selection method is compatible with alternating between positive and negative selections against different forms of amyloidogenic proteins via FACS, as we demonstrate using positive selections against QD-fibril conjugates and negative selections against the corresponding monomeric protein in solution. The technique could be readily extended to incorporate positive and negative selections against different forms of aggregates. For instance, antibodies isolated from immunization have previously been reported to selectively recognize a particular aggregate conformation present in one disease (e.g., Multiple Systems Atrophy or MSA) but not in a different aggregated conformation of the same protein in a related disease (e.g., Parkinson’s disease)^3^. Our approach could be used to isolate disease-specific conformational antibodies by alternating between rounds of positive and negative selections against different aggregated conformers (“strains”) of the same amyloid protein. Finally, the high-quality antibodies generated in this work against multiple complex antigens suggests our methodology could be readily extended to other types of insoluble antigens, including other aggregated proteins and potentially even membrane proteins.

## MATERIALS AND METHODS

### Preparation of antibody-quantum dot conjugates

Site-click Qdot 655 antibody labeling kit (Invitrogen, S10453) was used to separately conjugate an anti-tau antibody (Tau-5, AB_2721194) and anti-α-synuclein antibody (MJFR1, Abcam, ab138501) to dibenzocyclooctyne (DIBO) modified QDs. Briefly, 125 µg of capture IgG (Tau-5 or MJFR1) was buffer exchanged into antibody preparation buffer provided in the kit to a final volume of ∼80 µL and then mixed with 10 µL of β-galactosidase for 4 h at 37 °C. Following the incubation, 80 µL of GalT enzyme together with UDP-GalNAz were added and the reaction was carried out in Tris buffer (pH 7.0) overnight at 30 °C to generate IgG with azide-functionalized glycans. Following extensive washing steps using Tris buffer (pH 7.0), the azide-modified IgG (in a final volume of 100 µL) was mixed with 50 µL of QD solution for overnight incubation at room temperature. The resultant antibody conjugates were further purified with a Nanosep centrifugal filter (MWCO, 300 kDa) and resuspended into 250 µL of Tris buffer (pH 7.0). This final solution was further centrifuged at 4000x g for 5 min, and the supernatant was collected and stored at 4 °C until further use.

### Library sorting to identify lead tau antibodies

A human non-immune single-chain (scFv) library^14^ was sorted first against anti-Myc antibody (Cell Signaling, 2276S) to select for full-length scFv antibodies. Yeast cells (2x10^9^) displaying antibodies were incubated with anti-Myc antibody (80 nM) in 1% milk in PBSB (PBS with 1 g/L BSA) in a volume of 5 mL at room temperature with end-over-end mixing for 3 h. Following primary incubation, the cells were washed once with 25 mL of ice-cold PBSB and incubated with 1000 µL of Protein G microbeads (Miltenyi, 130-071-101) in a final volume of 40 mL of PBSB with end-over-end mixing at 4 °C for 45 min. Next, the cells were washed once with ice-cold PBSB, and the beads and cells were passed through an LS column (Miltenyi, 130-042-401), and the column was washed once with cold PBSB. Yeast cells bound to the beads were eluted from the column and grown in 50 mL of SDCAA media [16.75 g/L sodium citrate trihydrate, 4 g/L citric acid, 5 g/L casamino acid, 6.7 g/L yeast nitrogen base (without amino acids) and 20 g/L dextrose]. Dilutions of the collected cell culture were plated on yeast dropout plates. The number of cells retained in this selection was determined to be ∼8x10^8^.

Tau aggregates (100 µg; Stressmarq, SPR-329) were first sonicated on ice for 5 mins (30 s on, 30 s off) in a volume of 500 µL HEPES buffer. Tosyl Dynabeads (∼8x10^7^ beads; Invitrogen, 14203) were washed 2x with HEPES using a DynaMag-2 magnet (Invitrogen, 12321D). Next, the beads and sonicated fibrils were incubated in a final volume of 1 mL of HEPES buffer with end-over-end mixing at room temperature for 2-3 d. The beads were then stored at 4 °C prior to use.

The scFv library previously enriched for full-length antibody expression was next sorted via MACS against tau aggregates. For sort 1, 10^9^ yeast cells were washed twice with PBSB. Next, ∼10^7^ tosyl beads coated with tau aggregates were blocked twice with 10 mM glycine (pH 7.4) by placing the beads in a tube on a DynaMag-2 magnet for 5 min for each blocking step. This blocking step was followed by one wash with PBSB. Yeast and beads were incubated in PBSB with 1% milk in a volume of 5 mL with end-over-end mixing at room temperature for 3 h. Post incubation, the beads were washed once with ice-cold PBSB by placing the tube on a DynaMag-15 magnet (Invitrogen, 12301D). Yeast bound to beads were recovered in 50 mL of SDCAA media for 2 d. Dilutions were plated post washing to estimate the numbers of cells retained during the selection. Sorts 2 and 3 were performed in the same way except that a starting number of 10^7^ yeast cells were used and the incubation with tau aggregate-coated beads was performed in a final volume of 1 mL.

In sort 4, a negative FACS selection against tau monomer was performed. Tau monomer (with a C-terminal His-tag) was expressed in pLysS cells. Following expression, monomer was purified by incubating with Ni-NTA agarose beads (Qiagen, 30230) and 1 mM NiSO_4_ overnight at 4 °C with mild agitation. Tau monomer was eluted at low pH, aliquoted, and stored at -80 °C until use. To promote the monomeric conformation, tau monomer was thawed and incubated overnight with 4 M GnHCl at 37 °C. The monomer was then incubated with 10 mM DTT at 100 °C for 10 min, buffer exchanged twice into 20 mM HEPES buffer, and allowed to refold at room temperature for 1 h prior to use. The concentration of tau monomer was determined using BCA assay.

For sorting, 10^7^ yeast cells were washed twice with PBSB and incubated with 10 nM of His-tagged tau monomer in a final volume of 1 mL in PBSB at room temperature for 3 h with end-over-end mixing, Next, the cells were washed once with ice-cold PBSB and incubated with 1000x dilution of mouse anti-Myc and chicken anti-His (Invitrogen, PA1-9531) antibodies on ice for 20 min. Next, the cells were washed once with ice-cold PBSB and then incubated with a 200x dilution of goat anti-mouse Alexa Fluor 488 (Invitrogen, A11001) and a 1000x dilution of donkey anti-chicken IgY F(ab)’_2_ Alexa Fluor 647 (Jackson Immunoresearch, 703-606-155) on ice for 4 min. Post-secondary incubation, the cells were washed once with ice-cold PBSB and sorted on Beckman Coulter MoFlo Astrios sorter. Yeast cells displaying antibodies and with minimal binding to tau monomer were collected.

In round 5, a positive FACS selection was performed against tau fibril-QD conjugates. First, 5 µg (2.5 µL at 2 µg/µL) of tau fibrils was diluted to a final volume of 200 µL in PBS and sonicated on ice for 5 min (30 s on, 30 s off). Next, the fibrils were vortexed briefly and moved to a new tube. QD conjugates (5 µL) were added to tau fibrils and incubated with end-over-end mixing at room temperature for 2-3 h (referred to as the high antigen sorting condition). In parallel, 5 µg of tau monomer was diluted in HEPES (20 mM, pH 7.4) to a volume of 200 µL and incubated with 5 µL QD conjugates in a similar manner as done for fibrils. Yeast cells (10^7^) were washed twice with PBSB and incubated with either tau fibril-QD conjugates or tau monomer-QD conjugates in a volume of 200 µL with a 1000x dilution of mouse anti-Myc antibody in 1% milk in PBSB at room temperature for 3 h with end-over-end mixing. Post incubation, the cells were washed once with ice-cold PBSB and incubated with a 200x dilution of goat anti-mouse IgG Alexa Fluor 488 on ice for 4 min. Next, the cells were washed once with ice-cold PBSB and sorted on Beckman Coulter MoFlo Astrios sorter. Gates were drawn to maximize conformational specificity by collecting tau fibril binders over tau monomer binders.

### Tau antibody affinity maturation

The lead tau antibody clone (ATA1) isolated via the first set of MACS and FACS sorts was cloned into a modified version of pCTCON2 yeast surface display plasmid and sequence confirmed. Next, soft randomization libraries were prepared with diversity focused in HCDR1, HCDR2, LCDR1, and LCDR3 at sites in ATA1 that are indicated in **Fig. S1**. These libraries sampled the wild-type residue at a frequency of ∼50% and all other amino acids at a total frequency of ∼50%. The four single CDR libraries were prepared as described previously^13, 36^, and ∼4.8x10^7^ (LCDR1), 15.7x10^7^ (LCDR3), 23.8x10^7^ (HCDR1) and 10.1x10^7^ (HCDR2) transformants were obtained. Next, one round of positive selection against tau aggregates by MACS was performed, as described above. Yeast cells (10^9^) were washed twice with PBSB. In parallel, ∼10^7^ tosyl beads coated with tau fibrils were first blocked twice with 10 mM glycine using a DynaMag-2 magnet for 5 min for each blocking step. This blocking was followed by a wash with PBSB. Next, the beads and yeast were incubated in 1% milk in PBSB in a final volume of 5 mL with end-over-end mixing for 3 h at room temperature. Following primary incubation, the beads were washed once with ice-cold PBSB and yeast bound to beads were recovered in 50 mL of SDCAA media. Dilutions were plated to estimate the number of cells retained during the positive selection.

In round 2, a positive FACS selection was performed against tau fibril-QD conjugates. The conjugates were prepared as described above except that the mass of tau fibrils was reduced (2 µg instead of 5 µg) and the volume of QD conjugates was reduced to 2 µL from 5 µL. This preparation of conjugates is referred to as the low antigen condition. The FACS selection was performed as described above. The binding of the parental antibody (ATA1) was also examined at the same condition to select for drawing gates that collect antibody-displaying yeast cells with stronger binding than observed for the parental antibody. Cells were collected from three libraries (HCDR1, HCDR2 and LCDR3) that showed the highest levels of binding and higher than that for the parental antibody.

Next, plasmids from the enriched HCDR1, HCDR2 and LCDR3 libraries were isolated using a yeast mini-prep kit (Zymo, D2004). DNA segments encoding each of the three CDRs were then amplified by PCR. Overlap PCR was used to assemble these DNA segments into complete scFvs. This overlap PCR reaction produced a shuffled library that combined the diversity from the three enriched libraries (HCDR1, HCDR2 and LCDR3). The new sub-library was screened against tau fibrils by MACS. Yeast cells (10^9^) were washed twice with PBSB. Tosyl beads coated with tau fibrils (∼10^7^ conjugated beads) were first blocked with 10 mM glycine (pH 7.4) and washed once with PBSB. The beads and yeast were incubated and sorted, as described above. In round 2, a positive FACS selection was performed against tau fibril-QD conjugates, as described above for the high antigen condition. In round 3, a negative FACS selection was performed against tau monomer (10 nM), as described above. In the final round of sorting, a positive FACS selection was performed against tau fibrils-QD conjugates using the low fibril condition, as described above.

### Library screening for isolating and affinity maturing α-synuclein antibodies

The screening for conformational antibodies that recognize α-synuclein fibrils was performed in a similar manner as described for the tau antibodies. α-synuclein fibrils (100 μg; Stressmarq, SPR-317) were sonicated on ice for 5 min (30 s on, 30 s off). Next, 8x10^7^ tosyl beads were washed twice with PBS by placing them on the magnet. The fibrils were added to beads and incubated with end-over-end mixing at room temperature for 2-3 d. Beads immobilized with fibrils were moved to 4 °C prior to use.

The same human scFv library previously enriched for full-length expression (as described above) was also used for selections against α-synuclein fibrils. In round 1, MACS was performed in the largely the same way as the tau selections. Yeast cells (10^9^) were incubated with fibril-coated beads (10^7^). In rounds 2-4, MACS was performed with a reduced number of cells (10^7^) without reducing the number of beads (10^7^). In round 5, a negative FACS selection was performed for a lack of binding to disaggregated α-synuclein monomer. Yeast cells (10^7^) were incubated with 10 nM biotinylated α-synuclein monomer and mouse anti-Myc antibody (1000x dilution) at room temperature for 3 h with end-over-end mixing. Post primary incubation, the cells were washed once with ice-cold PBSB and incubated with a 200x dilution of goat anti-mouse Alexa Fluor 488 and 1000x dilution of streptavidin Alexa Fluor 647 on ice for 4 min. After incubation with secondary reagents, cells were washed once with ice-cold PBSB and sorted on a Beckman Coulter MoFlo Astrios sorter. Yeast cells that displayed antibodies and showed minimal binding to α-synuclein monomer were collected via FACS. In rounds 6-8, positive FACS selections were performed against 1x α-synuclein fibril-QD conjugates (high antigen condition). The selections were performed as described above for tau fibril-QD conjugates.

The lead α-synuclein antibody aS2.1 identified during initial discovery was cloned into a modified form of the pCTCON2 yeast-surface display plasmid in the format of a single-chain Fab (scFab). The antibody was cloned as an N-terminal fusion to Aga2 (V_L_-C_L_-linker-V_H_-C_H_1-Aga2) and sequence confirmed. Next, an HCDR2 library was prepared using soft randomization for affinity maturation by randomizing the sites in aS2.1 indicated in **Fig. S6**. The library was prepared as described previously^13, 36^. All selections were performed as described above. In round 1, MACS was performed with 10^9^ cells and 10^7^ beads. In rounds 2, 4 and 6, FACS positive selections were performed against α-synuclein fibril-QD conjugates (high antigen condition). In rounds 3 and 5, FACS negative selections were performed against a polyspecificity reagent (soluble membrane proteins in round 3), prepared as described previously^20, 41^, and disaggregated α-synuclein monomer (round 5).

### Antibody cloning and production

Antibody genes were isolated from the terminal rounds of sorting against tau and α-synuclein fibrils using a yeast miniprep kit (Zymo, D2004). For tau, scFv genes were amplified by PCR and ligated into a scFv-Fc mammalian expression plasmid (pTT5) using *NheI* and *HindIII* restriction sites. For α-synuclein, V_H_ genes were amplified by PCR and cloned into an IgG heavy chain expression plasmid (pTT5) using restriction sites *EcoRI* and *NheI*, and the V_L_ genes were amplified by PCR and cloned into an IgG light chain expression plasmid (pTT5) using restriction sites *EcoRI* and *BsiWI*. The ligation mixtures were next transformed into competent DH5α cells, plated on LB-agar plates (supplemented with 100 µg/mL ampicillin) and incubation at 37 °C for 16-20 h. The following day, individual colonies were picked from these plates, grown overnight in LB media (supplemented with 100 µg/mL of ampicillin), mini-prepped (Qiagen, 27106) and sequenced.

Antibodies were expressed with HEK293-6E cells (National Research Council of Canada). Cells were maintained and passaged in F17 media (Invitrogen, A1383502) supplemented with 0.1% w/v kolliphor (Fisher, NC0917244), 30 mL/L L-glutamine (Gibco, 25030081) and 0.5 mL/L geneticin (Gibco, 10131035) at a density of ∼1.7-2 million cells/mL, as described previously^42^. Cells were transfected at a density of ∼1.7-2 million cells/mL in F17 media (un-supplemented) with 15 µg of plasmid (15 µg for tau scFv-Fc plasmid or 7.5 µg each for a-synuclein IgG heavy and light chain plasmids) and 45 µg of PEI. Yeastolate (BD Sciences, 292804) was added at 20% w/v 24-48 h post transfection and cells were allowed to grow for additional 3-4 d. Post expression, cells were spun down at 3500xg for 40 min and media was moved to a new tube. Protein A agarose beads (300-500 µL of slurry; Fisher, 20334) were added to the tubes followed by gentle shaking overnight at 4 °C. The following day, the beads were collected in filter columns (Fisher, 89898) under vacuum and washed with 50 mL of PBS. Beads were next incubated with 1 mL 0.1 M glycine (pH 3.0) for 15 min to elute protein, and the eluate was collected by centrifugation. Proteins were next buffer exchanged into 20 mM acetate (pH 5.0) with Zeba desalting columns (Fisher, 89890), passed thorough 0.22 µm filters (EMD Millipore, SLGV004SL), aliquoted, and stored at -80 ᵒC. Protein concentrations were evaluated by measuring the absorbance at 280 nm, and the purity was analyzed by SDS-PAGE (Invitrogen, WG1203BOX).

### Flow cytometry analysis of antibody affinity and conformational specificity

Antibody affinity was analyzed via flow cytometry using a bead-based assay^13, 43^. Dynabeads coated with tau or α-synuclein aggregates (prepared as described above) were blocked with 10 mM glycine (pH 7.4) at room temperature for 1 h with end-over-end mixing followed by one wash with PBSB. Antibodies were thawed at room temperature, centrifuged at 21000xg for 5 min, and the supernatant was moved to a new tube. Antibody concentrations were next evaluated by measuring the absorbance at 280 nm. In each well of a 96-well plate (Greiner, 650261), 10^5^ fibril-coated beads were incubated with varying concentration of antibodies in PBSB supplemented with 1% milk for 3 h at room temperature with mild agitation. Post primary incubation, the beads were washed once with ice-cold PBSB and incubated with a 300x dilution of goat anti-human Fc, Alexa Fluor 647 (Jackson Immunoresearch, 109-605-098) on ice for 4 min. The beads were washed one more with ice-cold PBSB and analyzed on a Bio-Rad ZE5 flow cytometer, and the mean fluorescent values were measured.

Antibody conformational specificity was measured using a related bead-based assay^13, 43^. First, antibodies at fixed concentration (10 nM) were incubated with varying concentrations (0.1-1000 nM) of disaggregated tau or α-synuclein monomer in PBSB supplemented with 1% milk in a 96-well plate for 1 h at room temperature with mild agitation. Next, 10^5^ tau or α-synuclein aggregate-coated DynaBeads beads, which were blocked with 10 mM glycine as described above, were added to the antibody-monomer mixtures, and beads were allowed to incubate with the antibody-monomer mixtures for an additional 3 h. Post primary incubation, the beads were washed once with ice-cold PBSB and incubated with a 300x dilution of goat anti-human Fc Alexa Fluor 647 on ice for 4 min. The beads were washed once more and analyzed on Bio-Rad ZE5 flow cytometer. The median (tau) or mean (α-synuclein) fluorescent measurements of antibodies binding to fibrils were recorded.

### Preparation of recombinant tau proteins

The full-length human tau protein construct (2N4R/HT40) containing a C-terminal 6xHis tag in the pT7 plasmid was used to produce recombinant tau protein as previously described^4^. Site-directed mutagenesis (Agilent QuikChange Lightning, Santa Clara, CA, 210519-5) was used to introduce the following mutations in the HT40 construct: E3A, P4A, R5A, Q6A, E7A, FE8-9AA, V10A, M11A, E12A, D13A/H14A/A15A/G16A/T17A (X13-17A), Y18E, Y18F, Y18A/G19A/L20A/G21A (X18-21A) or D22A/R23A/K24A (X22-24A). All other Tau constructs (HT39, mouse tau, deletion constructs) are described elsewhere^4, 44^. These constructs were transformed into T7 Express competent cells (New England Biosciences, Ipswich, MA, C2566I). 125 mL of Luria Broth was inoculated with the bacteria and grown overnight at 25 °C until it reached an OD600 value of 0.8–1.0. IPTG was added to the flask at a final concentration of 1 mM and incubated at 37 °C for 2 h. The cells were centrifuged at 8000 ×g for 10 min, washed with saline buffer and centrifuged again at 8000 xg for 10 min. The pellet was frozen at -80 °C. The cells were resuspended in buffer (50 mM NaCl, 1 mM Tris, 500 nM imidazole and protease inhibitors at 10 μg/mL of pepstatin, leupeptin, bestatin, and aprotinin, and 1 mM PMSF, pH 8.0) and lysed by sonication (20 pulses of 5 s each at power setting of 3, Misonix XL-2000) on ice. Protease inhibitors were again added to the lysate at the same concentrations as above along with Brij 35 to 0.1%. The sample was then centrifuged at 107,000 ×g for 20 min at 4 °C. Protease inhibitors and 10% glycerol were added to the supernatant before loading it onto 1 mL of Talon metal affinity resin (Clontech, Mountain View, CA, #635502). After a 20 mL buffer wash, the protein was eluted into 1 mL fractions by adding 10 mL of elution buffer (50 mM NaCl, 1 mM Tris, and 100 mM imidazole with 2 mM PMSF). The samples were analyzed by SDS-PAGE and Coomassie staining and fractions with protein were combined and dithiothreitol (DTT) was added to 1 mM final concentration.

Protein concentrations were determined by SDS Lowry (Fisher, 23240) where 200 μL of the proteins were added to 200 μL of SDS solution (2% SDS, 5% β-mercaptoethanol, 10% glycerol, 62.5 mM Tris, pH 6.8), respectively, along with buffer-only blank samples. Protein standards of 0, 2, 4, 8, 16, and 32 μg were generated using a 2 mg/mL bovine serum albumin (BSA) stock (Thermo Fisher Scientific, Waltham, MA, #23209) and adding SDS solution to a total volume of 100 μL. One mL of 10% perchloric acid/1% phosphotungstic acid was added to each sample, vortexed, and briefly spun in microcentrifuge before a 1 h incubation on ice. The samples were then centrifuged at 18,000 xg for 15 min at 4 °C and the supernatant was removed. After the protein pellet had completely dried, it was dissolved in 1 mL of Lowry solution (0.01% CuSO_4_, 0.02% sodium potassium tartrate, and 2% sodium carbonate in 0.1 N sodium hydroxide) and incubated at room temperature for 10 min. 100 μL of Folin–Ciocalteu’s phenol reagent (diluted 1:1 in water, Sigma–Aldrich, St. Louis, MO, F9252) was added and incubated for 30-45 min at room temperature. 100 μL of each sample was added, in duplicate, to a clear 96-well plate and the absorbances were read at 750 nm. The total protein concentrations were interpolated from the BSA standard curve. Proteins were aliquoted and frozen at -80°C.

### Preparation of recombinant tau aggregates

Recombinant tau monomer and aggregate samples used in **Fig. S3** were generated by making a 400 μL reaction with 4 μM hT40 tau in polymerization buffer (5 mM DTT, 100 mM NaCl, 100 μM EDTA, and 10 mM HEPES at pH 7.6). The reaction was split in half and 7.5 μL of 2 mM arachidonic acid in ethanol was added to a final concentration of 75 μM in order to induce tau polymerization in the aggregate sample. The monomer sample received 7.5 μL of ethanol as a control. The aggregate reaction was allowed to proceed overnight. Aliquots of the samples were stored at -80°C until used.

The human tau-441 isoform (2N4R) used in **Fig. S18** was expressed recombinant in E. coli BL21 (DE3) cells and purified as described previously^54, 55^. To generate tau oligomers, the monomeric tau (1 mg/mL) was dialyzed against 1x PBS and stirred for 48 h at 37°C. The samples were aliquoted and stored at -80 °C until further use.

### Preparation of recombinant α-synuclein aggregates

Preformed fibrils of α-synuclein used in **Fig. S13** were generated as previously described^45^. Briefly, recombinant α-synuclein monomers for the generation of fibrils (Proteos, Kalamazoo, MI #RP-003) were diluted with dPBS to a final concentration of 5 mg/mL. The sealed tube of monomers was placed in an orbital thermoshaker for 7 d at 37 °C, shaking at 1,000 rpm. Aggregated α-synuclein was aliquoted and stored at -80°C until used.

Human α-synuclein used in **Fig. S18** was expressed recombinantly as described previously^55, 56^. Monomeric α-synuclein was stirred for 48 h at 37°C to produced oligomeric α-synuclein. The protein was aliquoted and stored at -80 °C until further use. To generate α-synuclein fibrils, samples were stirred for 6-7 d at 37 °C. Sodium azide was added at 0.01% to the final solution to avoid potential bacterial growth^57^.

### Preparation of Aβ oligomers

Aβ42 oligomers were prepared as described previously^58, 59^. Briefly, 1 mg Aβ42 pellet was dissolved in 600 µL of hexafluroisopropanol and incubated for 10-20 min at room temperature. The resulting solution was added to 2100 µL of molecular grade water in a siliconized Eppendorf tube with perforated caps to allow slow evaporation of hexafluroisopropanol. The samples were then stirred at 500 rpm using a Teflon-coated micro-stirred bar for 48 h at room temperature in the fume hood.

### Antibody epitope analysis using indirect ELISAs

Indirect ELISAs, as reported in **Fig. S5**, were performed to determine the binding epitope of ATA1.459.3 scFv-Fc fusion protein as previously described^48^. 50 μL of 2 ng/μL recombinant protein (human HT40, mouse HT40, and a series of overlapping deletion human tau protein constructs Δ2-18, Δ9-155, Δ50-244, Δ144-273, Δ209-356, Δ283-441, Nterm, NtermMTBR; numbering based on human 2N4R isoform of 441 amino acids) in borate saline solution (100 mM boric acid, 25 mM sodium tetraborate decahydrate, 75 mM NaCl, 250 μM thimerosal) was used to coat each well (Corning, 3590) for 1 h. Washing steps were done with ELISA wash solution (100 mM boric acid, 25 mM sodium tetraborate decahydrate, 75 mM NaCl, 250 μM thimerosal, 0.4% BSA and 0.1% Tween 20, pH 9.0) (200 μL per well). Wells were blocked with 5% non-fat dry milk made in ELISA wash solution for 1 h. ATA1.459.3 was diluted in blocking buffer at 1:10,000 (511 pM) and incubated for 2 h. Tau-13 (AB_2721193) and Tau-7 (AB_2721195) antibodies were diluted in blocking buffer at 1:10,000 (66667 pM) and used as a positive control for tau signals. Peroxidase-conjugated goat anti-human IgG Fcγ fragment (Jackson ImmunoResearch, 109-035-190) was added at 1:5,000 dilution for 1 h. Reactivity was detected by adding 3,3′,5,5′ tetramethylbenzidine substrate (Sigma, T0440) and incubating for 2 min. Reactions were quenched with 50 μL of 3.6% H_2_SO_4_ and absorbance read at 450 nm. Absorbance (A) is not linear [i.e., A = Log10(1/transmittance)], thus, the absorbance data were converted to percent absorbed light (a linear scale) using the following equation %A = (1 – 10–x)∗100, where x is absorbance.

### Antibody epitope analysis using peptide scanning

Peptide scanning of the epitope of ATA1.459.3 was first performed using overlapping 15 amino acid peptides scanning the full-length tau sequence synthesized on PepSpots cellulose membranes (JPT Peptide Technologies). Neighboring tau peptides had 11 overlapping amino acids. The peptide membranes were rinsed with methanol for 5 min and washed with TBST (50 mM Tris, 137 mM NaCl, 2.7 mM KCl, 0.05% Tween-20, pH 8.0) three times for 3 min. Membranes were blocked with 5% milk in TBST at room temperature for 1 h. The membrane containing tau peptides was incubated with 0.01 µg/mL ATA1.459.3 (87.0 pM) in 5% milk in TBST at room temperature for 2 h. Membranes were then washed with TBST three times for 5 min. Membranes were incubated with HRP-conjugated goat anti-human IgG (1:5000) in 5% milk in TBST at room temperature for 1 h and were washed with TBST three times for 5 min. Membranes were incubated with SuperSignal West Femto Maximum Sensitivity Substrate (Thermo Scientific) for 1 min and imaged using the ChemiDoc MP imaging system (Bio-Rad).

To characterize the binding epitopes of ATA1.459.3 with greater precision, additional PepSpots membranes were manufactured with peptides based on information obtained from the initial round of epitope mapping. Epitope mapping with these membranes was performed as described above except 0.005 µg/mL ATA1.459.3 (43.5 pM) was used during the primary incubation.

### Sandwich ELISA analysis of tau and α-synuclein conformational specificity

Recombinant tau and α-synuclein sandwich ELISAs, as reported in **Fig. S3** for tau and **Fig. S13** for α-synuclein, were used to measure the differential reactivity of ATA1.459.3 (scFv-Fc fusion), zagotenemab, and Tau-5 for monomeric and aggregated tau samples and H2.7, cinpanemab, and LB509 (Abcam, Waltham, MA, #ab27766) for monomeric and aggregated α-synuclein samples in non-denaturing conditions. Capture antibodies were diluted to 2 ng/μL and 50 μL was added per well in a 96-well plate and allowed to incubate for 1 h. All incubations were carried out at room temperature and on a shaker. The wells were washed twice in ELISA wash buffer. The wells were blocked with 200 μL of wash buffer containing 5% non-fat dry milk for 1 h and then rinsed twice with wash buffer. The monomer and aggregated samples were serially diluted in tris buffered saline (TBS, 150 mM NaCl, 50 mM Tris, pH 7.4) from 240 nM to 12.2 pM at 1:3 intervals and 50 μL was added to the wells for 90 min then washed four times with wash buffer. The detection antibody (R1, a pan-tau polyclonal rabbit antibody^49^ for tau (AB_2832929); EPR20535 (Abcam, Waltham, MA, #ab212184), a monoclonal rabbit antibody for α-synuclein) was diluted at 1:10,000 (R1) or 1:1,000 (EPR20535) in 5% non-fat dry milk and 50 μL was added to the plates for 90 min. The wells were washed four times and 50 μL HRP-conjugated goat anti-rabbit (Vector Laboratories, PI-2000) detection secondary antibody was added at a 1:5000 dilution in milk for 1 h. Then the wells were washed four times in wash buffer followed by development with 50 μL of TMB for 5-20 min, stopped with addition of 50 μL of 3.6% H_2_SO_4_, and absorbance was read at 450 nm and converted to %A as above.

Sandwich ELISAs, as reported in **Fig. S8** for tau, were repeated as above, except where noted, with soluble tau fractions from control and AD human frontal cortex lysate samples. For the soluble tau samples, 50 μL containing 20 μg of total protein (0.4 mg/mL; diluted in tris buffered saline) was added to each well. The R1 tau antibody and HRP-conjugated anti-rabbit antibodies were used as above. The ELISAs were developed for 5-7 min.

### Immunoblotting and western blotting of mouse brain samples

This study was conducted in a facility approved by the American Association for the Accreditation of Laboratory Animal Care, and all experiments were performed in accordance with the National Institutes of Health Guide for the Care and Use of Laboratory Animals and approved by the Institutional Animal Care and Use Committee of the University of Michigan. Mice were housed at the University of Michigan animal care facility and maintained according to U.S. Department of Agriculture standards (12 h light/dark cycle with food and water available ad libitum). Hemizygous P301S tau mice (B6;C3-Tg-Prnp-MAPT-P301S PS19Vle/J; The Jackson laboratory stock #008169)^51^, homozygous A53T mice (B6;C3-Tg(Prnp-SNCA*A53T)83Vle/J; The Jackson laboratory stock #004479)^52^ and non-transgenic littermates were bred at the University of Michigan and euthanized at 9, 10 and 11 months for brain collection.

Animals were deeply anesthetized with isofluorane and perfused transcardially with 1x PBS. Brains were divided sagittally. One half was immediately placed on dry ice and stored at -80 °C for biochemical studies while the other half was fixed in 4% paraformaldehyde at 4 °C for 24 h, and cryoprotected in a series of 10% and 30% sucrose in 1x PBS at 4 °C until saturated. Fixed hemispheres were snap frozen in OCT medium and sectioned at 12 μm sagittally using a cryostat and sections were stored at -20 °C for immunofluorescence.

The 11-month-old P301S, 10-month-old A53T and non-transgenic littermate brain samples were homogenized in PBS with a protease inhibitor cocktail (Sigma Aldrich, 11873580001) using a 1:3 dilution of tissue:PBS (w/v). Samples were centrifuged at 9300xg for 10 min at 4 °C. Supernatants (soluble fraction) were snap frozen and stored at -80 ℃ for Western blot analysis. Pellets were resuspended in PBS with protease inhibitor cocktail (Roche, 11836170001), centrifuged at 9300xg for 10 min (4 °C), and supernatants were discarded. The pellet was resuspended in 1% sarkosyl with protease inhibitor, vortexed (1 min), and incubated at room temperature (1 h). Samples were sonicated (water bath sonicator) for 5 min and centrifuged for 30 min (16000xg at 4 °C). Sarkosyl (insoluble) fractions of brain extracts (7 µg of total protein) were spotted directly onto nitrocellulose membranes and allowed to dry (1 h). Control dot blots (loading controls) were stained with Ponceau S (5 min) and washed 3x with distilled water. The other dot blots were blocked with 10% nonfat dry milk in Tris Buffered Saline with 0.1% Tween-20 (TBST) buffer at room temperature (1 h).

Tau dot blots, as reported in **Fig. S6A**, were next incubated with antibodies at 10 nM (1% nonfat dry milk in TBST) overnight at 4 °C. α-synuclein dot blots, as reported in **Fig. S14**, were incubated with 50 nM antibody (aS2.1 WT, H2.7, H2.4, H2.3) or a 1000x dilution of 5C2 (Novus Biologicals, NBP1-04321), LB509 (Abcam, ab27766) and MJFR1 (Abcam, ab2099538). Next, the blots were washed with TBST and incubated with a 5000x diluted solution of HRP-conjugated goat anti-human IgG or a 1000x diluted solution of HRP-conjugated goat anti-mouse or goat anti-rabbit at room temperature for 1 h. Afterward, the blots were washed with TBST and developed using Ecobright Pico HRP Substrate (Innovative Solutions) and visualized with the Genesys G:Box imaging system (Syngene). Three independent repeats were performed.

For western blotting reported in **Fig. S6B**, 25 µg of total protein was loaded on precast NuPAGE 4-12% Bis-Tris gels (Invitrogen). Gels were subsequently transferred onto nitrocellulose membranes and first stained with Ponceau S and washed 3x with distilled water. After imaging, membranes were destained for 1 min with 0.1 M NaOH and washed 3x with distilled water. Next, membranes were blocked for 1 h at room temperature with 10% nonfat dry milk in TBST buffer. Membranes were probed overnight at 4 ℃ with anti-tau and α-synuclein antibodies (10 nM) in TBST with 1% milk. HRP-conjugated goat anti-human IgG (5000x dilution) was used for detection. Ecobright Nano HRP Substrate (Innovative Solutions) was used to visualize bands with the Genesys G:Box imaging system (Syngene). Three independent repeats were performed.

### Immunofluorescent staining of mouse brain samples

Fixed 9-month-old P301S brain sections were post-fixed for 10 min in methanol at 4 ℃. Sections, as analyzed in **Fig. S7**, were washed in 1x PBS three times for 10 min and subjected to heat-induced antigen retrieval in 10 mM Citrate Buffer (pH 6). Sections were washed in 1x PBS two times for 5 min and permeabilized with 0.5% Triton-X 100, washed for 10 min in 1x PBS, and blocked using the Mouse on Mouse (M.O.M.) Blocking Reagent (M.O.M. Immunodetection Kit, Vector, BMK-2202) for 1 h. Sections were washed two times for two min in 1x PBS and incubated for 5 min in M.O.M. diluent. Sections were then incubated in ATA1.459.3 (100 nM) and AT8 (1:200, anti-phosphorylated tau, pS202 and pT205; Invitrogen) in M.O.M. diluent overnight at 4 ℃. The following day, sections were washed in 1x PBS three times for 10 min each and incubated with goat anti-mouse IgG Alexa Fluor 488 (Invitrogen; 1:500) and anti-human IgG Alexa Fluor 647 (1:500) for 1 h. Sections were then washed in PBS three times for 10 min each and incubated with DAPI (Sigma) to label nuclei for 5 min at room temperature, washed three times for 5 min each, and were mounted with Prolong Gold Antifade Reagent (Invitrogen). Slides were imaged using a Leica SP5 Confocal microscope.

### Immunofluorescent staining of human brain samples

Paraffin-embedded brain tissue sections from the midbrain was attained from subjects with Parkinson’s disease, dementia with Lewy bodies, and age-matched control subjects, as well as mid-frontal cortex from progressive supranuclear palsy, Alzheimer’s disease and age-matched control subjects from the Michigan Brain Bank (University of Michigan, Ann Arbor, MI, USA). Brain tissue was collected with patient consent and protocols were approved by the Institutional Review Board of the University of Michigan and abide by the Declaration of Helsinki principles. Samples were examined at autopsy by neuropathologists for diagnosis.

Paraffin sections from human brains were heated, deparaffinized and rehydrated with sequential dilutions of xylene, ethanol and distilled water. Sections were subjected to heated antigen retrieval in 10 mM Citrate Buffer (pH 6) and permeabilized with 0.5% Triton-X 100. Sections were incubated in 70% ethanol for 5 min and then incubated in autofluorescence eliminator reagent (Millipore catalog #2160) for 5 min and washed three times in 70% ethanol. Sections were blocked using 5% normal goat serum in 1x PBS for 1 h at room temperature. Tauopathy sections, as reported in **Fig. 4**, were incubated with either Tau-5 (Biolegend; 1:200) or AT8 (Invitrogen, 1:200) and ATA1.459.3 (100 nM). Synucleinopathy sections, as reported in **Fig. 6**, were incubated with either LB509 (Abcam, ab27766, 1:200) or EP1536Y (pS129, Abcam, ab51253, 1:500) and H2.7 (100 nM) overnight at 4 ℃. The following day, sections were washed in PBS three times for 10 min each and incubated for 1 h with goat anti-mouse or rabbit IgG Alexa Fluor 488 and goat anti-human IgG Alexa Fluor 647, respectively (Invitrogen; 1:500). Sections were then washed in PBS three times for 10 min each and incubated with DAPI (Sigma) for 5 min at room temperature to label nuclei, washed three times for 5 min each, and coverslipped with Prolong Gold Antifade Reagent (Invitrogen). Slides were imaged with a Leica SP5 confocal microscope.

### Immunohistochemical staining of human brain samples

Paraffin-embedded brain tissue, as analyzed in **Fig. 7**, was obtained from a Michigan Brain Bank as described above. Tissue was obtained from the frontal cortex of a sample with a high level of Alzheimer’s disease neuropathological change, and this tissue was determined by neuropathological diagnosis to contain both aggregated tau and aggregated α-synuclein pathology. Control tissue was obtained from the frontal cortex of a sample determined by neuropathological diagnosis to lack both aggregated tau and aggregated α-synuclein. Samples were transferred to the University of Michigan Rogel Cancer Center Tissue and Molecular Pathology Shared Resource (TMPSR) Core for immunohistochemical staining. Staining was performed using a DAKO Autostainer Link 48 (Agilent, Carpiteria, CA). For detection of antibodies with human IgG1 Fc (ATA1.459.3, zagotenemab, H2.7, and cinpanemab), a Human-on-Human HRP-Polymer kit (Biocore Medical, BRR4056KG) was used, and antibodies with human Fc were tagged with Digoxigenin for detection. For detection of antibodies with mouse Fc (Tau-5 and LB509), a goat anti-mouse Flex HRP detection antibody (Agilent, Carpiteria, CA) was used. All tissue staining was performed at room temperature. Brain sections were first deparaffinized in xylene, rehydrated using graded alcohols to water, and then rinsed in TBST. A Dako Envision Flex Target Retrieval Solution, Low pH was then used to perform heat-induced epitope retrieval for all antibodies except for LB509 that required no pretreatment. The slides were then blocked for 5 min using peroxidase block. Primary antibodies (ATA1.459.3, zagotenemab, Tau-5, H2.7, cinpanemab, and LB509) were then applied to slides containing consecutive brain sections. Primary antibody incubation was carried out for 1 h for antibodies with human Fc and for 30 in for antibodies with mouse Fc. Mouse anti-Digoxigenin secondary antibody was applied to the slides for the detection of Digoxigenin-tagged antibodies with human Fc for 15 min. Goat anti-mouse secondary antibody was applied to the slides for the detection of antibodies with mouse Fc for 20 min. The slides were then rinsed with TBST. Next, the slides stained with antibodies with human Fc (ATA1.459.3, zagotenemab, H2.7, and cinpanemab) were incubated with MACH 2 mouse HRP polymer for 30 min. All slides were then rinsed once with TBST, and then incubated with 3,3’diaminobenzidine (DAB) for 10 min. The slides were then rinsed once with DI water and counterstained with hematoxylin for nuclei detection. The slides were then rinsed with DI water, dehydrated using graded alcohols, and then cleared using xylene and coverslipped. After staining, the slides were submitted to the Digital Pathology slide scanning service, part of the University of Michigan Department of Pathology, Michigan Medicine, which provided assistance with the generation of whole-slide images.

Inferior temporal gyrus sections from control (Braak I–II, n = 3) and severe AD (Braak V–VI, n = 3) obtained from the Brain Bank of the Cognitive Neurology and Alzheimer’s Disease Center at Northwestern University (see **Table S1** for details on human subjects) and Parkinson’s disease (PD) substantia nigra sections (n=3) and control substantia nigra sections (n=2) from the Alzheimer’s Disease Research Center at Banner Sun Health Research Institute^46^ were used to evaluate the pattern of immunohistochemistry (IHC) staining with ATA1.459.3 and H2.7, as reported in **Fig. S16**, using established methods^7, 50^. Wash steps were performed 6× for 10 min in 0.1% Triton-X/TBS between all steps in the procedure. The tissue was washed and then incubated in 3% hydrogen peroxide to quench endogenous peroxidase activity for 1 h at room temperature. The tissue was then blocked in blocking buffer (10% goat serum/2% BSA/0.4% Triton-X in TBS) followed by incubation in the primary antibody (diluted in 2% goat serum/0.1% Triton-X/TBS) overnight at 4°C. The following dilutions were used of biotinylated primary antibodies: ATA1.459.3 1:200, H2.7 1:400. Initial titering experiments ensured that antibodies were used at optimal dilutions. The following day, the tissue was incubated in ABC solution (1 drop of solutions A and B/10 mL, Vector Laboratories, PK-6100) for 1 h at room temperature followed by development in 0.05% 3,3′-diaminobenzidine (DAB), 0.003% hydrogen peroxide, 0.1% Triton-X/TBS for 10 min. After development, the tissue was mounted and coverslipped with Cytoseal 60 mounting media (Thermo Fisher Scientific, #8310-16). Tissue sections from each case were processed simultaneously for each antibody staining to reduce variability between staining runs.

### Tau extraction from human frontal cortex tissue

Fresh frozen tissue samples from the frontal cortex of age-matched non-demented (control, n = 4), severe Alzheimer’s Disease (AD) cases (n = 4), and progressive supranuclear palsy (PSP) cases (n=4) were obtained from the Brain Bank of the Cognitive Neurology and Alzheimer’s Disease Center at Northwestern University, Alzheimer’s Disease Research Center at Banner Sun Health Research Institute^46^, and the University of Michigan Brain Bank (see **Table S1** for details on human subjects) and used for analysis in **Fig. S9C-D**. Soluble tau fractions were obtained as described previously^47^. The tissue was suspended in 10 volumes of Brain Homogenization Buffer (50 mM Tris, pH 7.4, 274 mM NaCl, 5 mM KCl with pepstatin, leupeptin, bestatin, and aprotinin protease inhibitors at 10 μg/mL and PMSF at 1 mM). The tissue was homogenized with a Tissue Tearor Model 985-370 on setting #3 for three times 10 s then centrifuged at 27,000 × g for 24 min at 4 °C. The supernatant fraction was collected. A Bradford protein assay using Bio-Rad Protein Assay Dye Reagent Concentrate (Bio-Rad, Hercules, CA, #5000006) was performed on the soluble tau fractions to determine total protein concentrations. The fractions were aliquoted and stored at -80 °C until used in assays outlined below.

### Western blotting analysis of tau and α-synuclein antibodies

For western blotting analysis of tau in **Fig. S3D-G**, 375 ng of the purified recombinant tau protein monomer and aggregate samples were loaded per lane. For similar analysis of α-synuclein in **Fig. S13D-F**, 304 ng of α-synuclein protein monomer and aggregate were loaded per lane. Brain lysates were loaded at 30 μg/lane of total protein. After a blocking step of 1 h with 2% milk/TBS, the membranes were probed overnight at 4 °C with primary antibodies. Tau blots were probed with ATA1.459.3 (1:1,000) and Tau-5 (1:100,000) as a loading control to normalize tau signals, then reprobed with GAPDH (1:2000, Cell Signaling D16H11 XP Rb, Danvers, MA) to serve as a loading control. α-synuclein blots were probed with H2.7 (1:1,000) and EPR20535 (1:1,000) as a loading control to normalize α-synuclein signals, and then reprobed with GAPDH (1:2000) to serve as a loading control. Blots were rinsed with 0.1% Tween 20/TBS and then incubated with appropriate secondary antibodies diluted 1:20,000 in milk/TBS (IRDye 680RD goat anti-mouse IgG1, #926-68050; IRDye 800CW goat anti-Mouse IgG1, #926-32350; and IRDye 800CW goat anti-rabbit IgG (H + L), #926-32211; all from Li-Cor Bisciences, Lincoln, NE; Alexa Fluor® 680 Goat Anti-Human IgG, Fcγ fragment from Jackson ImmunoLabs, West Grove, PA, # 109-625-190). Blots were imaged using a Li-Cor Odyssey infrared imaging system in the 700 and 800 nm channels at 42 μm resolution.

Frozen brain tissue from the substantia nigra was obtained from Lewy Body Dementia (LBD) and age-matched control subjects as well as cortex from Alzheimer’s Disease (AD) and cortex from age-matched control subjects from the Michigan Brain Bank (University of Michigan, Ann Arbor, MI) as described in **Table S2**. Brain tissue was collected with patient consent, and protocols were approved by the Institutional Review Board of the University of Michigan and abide by the Declaration of Helsinki principles. Brain samples were examined at autopsy by neuropathologists for pathologic diagnosis. Lysates were prepared as previously described^53^.

For western blotting of the anti-tau antibodies (zagotenemab and ATA1.459.3 scFv-Fc fusion), as shown in **Fig. S9A-B**, 15 µg of PBS-soluble protein from the lysates of the cortex was loaded onto precast NuPAGE 4-12% Bis-Tris gels (Invitrogen). For western blotting of the anti-α-synuclein antibodies (cinpanemab and H2.7), as shown in **Fig. S15A-B**, 30 µg of PBS-soluble protein from the lysates of the substantia nigra was loaded. Gels were subsequently transferred at 35 V for 16 h onto 0.22 µm nitrocellulose membranes and first stained with Ponceau S and washed 3x with distilled water. After imaging, membranes were washed 3x with distilled water. Next, membranes were blocked for 30 min at room temperature with 5% nonfat dry milk in TBST buffer. Membranes were probed overnight at 4 ℃ with anti-tau (5 nM) and anti-α-synuclein (10 nM) antibodies in TBST with 1% milk. HRP-conjugated goat anti-human IgG-Fc (10000x dilution) was used for detection (Invitrogen Cat# 31413). Immobilon Western Chemiluminescent HRP Substrate (Millipore Sigma) was used to visualize bands with the Genesys G:Box imaging system 3 (Syngene). Parallel blots of the tissue lysates were run and probed with secondary antibody only under the same conditions to account for any non-specific binding of human tissue to human secondary antibody.

For western blots reported in **Fig. S9C-D**, human frontal cortex lysates from controls (n = 4), AD (n = 4) and PSP (n=4) were precleared by incubating with Protein A Mag Sepharose (Cytiva, Marlborough, MA, #28-9440-06) for 45 min to minimize cross-reactivity from endogenous human IgG. Recombinant protein samples and precleared brain lysate samples were prepared in Laemmli buffer (final 1× composition: 20 mM Tris, pH 6.8, 2% SDS, 6% glycerol, 1% β-mercaptoethanol, 0.002% Bromophenol Blue). All samples were heated at 99 °C for 5 min and separated using Novex Precast 4–20% gels (Invitrogen, Waltham, MA, #WXP42020BOX) in SDS-PAGE. Proteins were then transferred to nitrocellulose filter paper (Invitrogen, Waltham, MA, #LC2009) at 400 mA for 50 min.

### Analytical size-exclusion chromatography

Antibodies were analyzed via size-exclusion chromatography (SEC) with a Shimadzu Prominence HPLC System. After purification by Protein A, antibodies were buffer exchanged into 1x PBS. 100 µL of 0.1 mg/mL antibody was then injected into a SEC column (Superdex 200 Increase 10/300 GL column; GE, 28990944) and analyzed at 0.75 mL/min using a 1x PBS running buffer with 200 mM arginine (pH 7.4). Absorbance was monitored at 280 nm, and the percentage of antibody monomer was calculated by analyzing absorbance peaks between the void volume and buffer elution times.

### Immunoblotting of tau and α-synuclein conformers

Immunoblots shown in **Fig. S17** were prepared by blotting tau and α-synuclein proteins onto nitrocellulose membrane (GE Healthcare Life Science, 10600004). Human HT40 fibrils (Stressmarq, SPR-329), human 0N3R fibrils (Stressmarq, SPR-491), human K18 P301L fibrils (Stressmaq, SPR-330), mouse 2N4R fibrils (Stressmarq, SPR-475), human α-synuclein fibrils (Stressmarq, SPR-317), human N-acetylated α-synuclein fibrils (Stressmarq, SPR-332), and mouse α-synuclein fibrils (Stressmarq, SPR-324) were first sonicated for 5 min (30 s on, 30 s off) before loading onto the nitrocellulose membrane. Human α-synuclein oligomers (Stressmarq, SPR-484) were loaded onto the nitrocellulose membrane without sonication. Equal mass (50 ng) of all proteins was loaded onto nitrocellulose membrane, and loading was confirmed by silver stain detection.

Immunoblotting was performed at room temperature with mild rocking. The membranes were first blocked with 10% milk for 1 h. The membranes were then washed three times with 1x PBS supplemented with 0.1% Tween 20 (Sigma Aldrich, P1379; PBST) for 5 min. The membranes were then incubated with primary antibodies in PBST supplemented with 1% milk for approximately 3 h. Tau antibodies (ATA1.459.3 and zagotenemab) were incubated with membranes at a concentration of 0.2 nM, and α-synuclein antibodies (H2.7 and cinpanemab) were incubated with the membranes at 1 nM. After the primary incubation, the membranes were washed three times with PBST. The membranes were then incubated with a 1:10,000 dilution of goat anti-human Fc HRP secondary antibody (Invitrogen, A18817) in PBST for 1 h. The membranes were then washed twice with PBST and twice with 1x PBS. Immunoblots were then developed with ECL Western Blotting Substrate (Pierce, 32109). Signals were evaluated using X-ray films (Thermo Scientific, 34091), and films were developed using an X-ray film developer. Immunoblotting was repeated three times, and a representative image is shown.

Dot blot assays to investigate the antibody affinities to α-synuclein and tau species, as reported in **Fig. S18**, were performed as described previously^60^. Briefly, the protein stocks at 1 mg/mL were used, and 1.2 µL of sample was applied on to nitrocellulose membranes and then blocked with 10% nonfat milk in Tris-buffered saline with 0.01% Tween (TBST) overnight at 4°C. Membranes were then probed with humanized ATA1.459.3, zagotenemab, H2.7, and cinpanemab (1:1000) diluted in 5% nonfat milk for 1h at room temperature. HRP-conjugated IgG anti-human (1:5,000) was used to detect the different antibody affinities with different α-synuclein and tau species.

### Polyspecificity analysis

The polyspecificity reagent (soluble membrane proteins, SMP) was prepared as previously described^20, 41^. For biotinylation, sulfo-NHS-LC-biotin (Thermo Fisher, PI21335) was dissolved in distilled water at ∼11.5 mg/mL. Stock solution of Sulfo-NHS-LC-biotin (150 µL) and the SMP reagent (4.5 mL at 1.0 mg/mL) were mixed end-over-end mixing at room temperature (45 min). The reaction was quenched (10 µL of 1.5 M hydroxylamine at pH 7.2), and biotinylated SMP was aliquoted and stored at -80 ᵒC. The specificity assay was performed as previously described^13, 20^. Three independent repeats were performed and were normalized between high (emibetuzumab) and low (elotuzumab) polyspecificity control mAbs at the highest evaluated antibody concentrations.

### Antibody melting temperature analysis

The stability of antibodies was evaluated by differential scanning fluorimetry. The antibodies were diluted to 0.12 mg/mL and then mixed with Protein Thermal Shift Dye (Applied Biosystems, 4461164) to achieve a final concentration of 1x in final volume of 20 µL. Samples were submitted to the University of Michigan Advanced Genomics Core for analysis. The measurements were taken on ABI Prism 7900HT Sequence Detection System (Applied Biosystems), and the temperature range examined spanned between 25 and 98 °C. Three independent repeats were performed for this study.

### Charged-stabilized self-interaction nanoparticle spectroscopy (CS-SINS)

CS-SINS was measured as reported previously^25^. Briefly, capture antibody (Jackson ImmunoResearch, 109-005-008) and polylysine (90%:10% w/w ratio respectively; Fisher Scientific, ICN19454405) were immobilized on concentrated gold nanoparticles and incubated overnight. Dilute mAb solutions (11.1 μg/mL, 45 μL) were incubated with 5 μL of gold conjugates for 4 h at room temperature. Absorbance spectra was measured on a Biotek Synergy Neo plate reader (Biotek, Winooski, VT) in 1 nm increments between 450 and 650 nm. The inflection of a quadratic equation fit describing the forty data points surrounding the maximum absorbance was calculated to determine the plasmon wavelength. Plasmon wavelengths were normalized between two parameters and calibrated against a panel of five antibodies (NIST, ibalizumab, mepolizumab, trastuzumab, and romosozumab). This calibration panel was used to rescale the measurements to the same scale as reported in the original study^25^.

## LIST OF ABBREVIATIONS

QD: quantum dot
FACS: fluorescence-activated cell sorting
MACS: magnetic-activated cell sorting
CDR: complementarity-determining region
Tm: melting temperature
CS-SINS: charge-stabilized self-interaction nanoparticle spectroscopy
IHC: immunohistochemistry

## DECLARATIONS

### Ethics approval and consent to participate

Brain tissue was collected with patient consent and protocols were approved by the Institutional Review Board of the University of Michigan and abide by the Declaration of Helsinki principles.

### Consent for publication

Not applicable.

### Availability of data and materials

The datasets used and/or analyzed during the current study are available from the corresponding author on reasonable request.

### Authors’ contributions

A.A.D., J.M.Z. and P.M.T. designed the research, A.A.D., J.H., and H.C. generated the QD-amyloid conjugates, A.A.D performed the antibody library sorting, A.A.D., J.M.Z., and J.H. produced the antibodies, A.A.D., J.M.Z, H.T., J.E.G, E.K.M., N.M.K. and K.N.D characterized the antibodies, H.T., J.E.G., H.P., N.M.K. and K.N.D performed and/or analyzed the immunostaining results, M.D.S., N.M.K. and K.N.D analyzed antibody binding data, and A.A.D, J.M.Z and P.M.T. wrote the paper with input from the co-authors. All authors read and approved the final manuscript.

## Acknowledgements

We thank K. Dane Wittrup for providing the library used for antibody selections. We thank Rakez Kayed for assistance with characterization of the antibodies reported here. We thank Kathy Toy from the University of Michigan Histology core for assistance with IHC staining. We thank Peter Ouillette from the University of Michigan Digital Pathology slide scanning service for assistance with imaging of IHC slides. We thank members of the Tessier lab for their helpful suggestions.

