## Supplemental Tables and Figures for "Flow cytometric isolation of drug-like conformational antibodies specific for amyloid fibrils"

**Table S1. Human subject data for samples isolated from human frontal cortex.** Tissues were obtained from the Brain Bank of the Cognitive Neurology and Alzheimer's Disease Center at Northwestern University, Alzheimer's Disease Research Center at Banner Sun Health Research Institute, and the University of Michigan Brain Bank. This tissue was used to prepare brain lysates analyzed in Figs. S8 and S9D-F and tissue sections analyzed via immunohistochemistry in Fig. S16.

| Diagnosis | Age (years) | Gender | Post-mortem interval (PMI) | Tau staging | Region | Sample type | Source |
|---|---|---|---|---|---|---|---|
| AD | 90 | M | 19.00 | Braak V | FtCtx | Frozen | NWU |
| AD | 87 | F | 3.00 | Braak V | FtCtx | Frozen | Banner |
| AD | 89 | F | 2.33 | Braak V | FtCtx | Frozen | Banner |
| AD | 60 | F | 3.50 | Braak VI | FtCtx | Frozen | Banner |
| AD | 84 | F | 6.00 | Braak VI | ITG | Fixed | NWU |
| AD | 91 | M | 4.00 | Braak VI | ITG | Fixed | NWU |
| AD | 86 | M | 10.00 | Braak VI | ITG | Fixed | NWU |
| ND | 78 | F | 2.83 | Braak III | SN | Fixed | Banner |
| ND | 61 | M | 2.33 | Braak I | SN | Fixed | Banner |
| ND | 88 | F | 3.00 | Braak II | FtCtx | Frozen | Banner |
| ND | 91 | M | 1.50 | Braak II | FtCtx | Frozen | Banner |
| ND | 80 | M | 3.25 | Braak II | FtCtx | Frozen | Banner |
| ND | 86 | M | 3.00 | Braak I | FtCtx | Frozen | Banner |
| ND | 53 | F | 11.00 | NA | FtCtx | Frozen | UofM |
| ND | 83 | F | Unknown | Braak I | ITG | Fixed | NWU |
| ND | 77 | M | 17.00 | Braak II | ITG | Fixed | NWU |
| ND | 82 | F | 5.00 | Braak II | ITG | Fixed | NWU |
| PD | 79 | M | 3.25 | Braak IV | SN | Fixed | Banner |
| PD | 85 | M | 2.16 | Braak IV | SN | Fixed | Banner |
| PD | 79 | M | 2.50 | Braak II | SN | Fixed | Banner |
| PSP | 87 | M | 23.00 | NA | FtCtx | Frozen | NWU |
| PSP | 68 | F | 11.00 | NA | FtCtx | Frozen | NWU |
| PSP | 66 | M | 5.00 | NA | FtCtx | Frozen | UofM |

**Table S2. Description of the human brain lysates from the Michigan Brain Bank.** These lysates were used for analysis in Figs. S9A-B and S15.

| **BBID** | **Age** | **Sex** | **Neuropathological diagnosis** |
|---|---|---|---|
| 2587 | 80 | M | High Likelihood AD |
| 2538 | 82 | F | High Likelihood AD |
| 2528 | 76 | F | High Likelihood AD |
| 1316 | 87 | M | Lewy body disease, diffuse neocortical subtype |
| 1417 | 72 | M | Lewy body disease, diffuse neocortical subtype |
| 681 | 84 | F | Lewy body disease, diffuse neocortical subtype |
| 1172 | 65 | M | Control |
| 619 | 83 | F | Control |
| 1874 | 71 | M | Control |

**ATA1.WT**

$V_H$ QVQLQQSGPGLLKPSETLSLTCVISGDSVSSNSATWNWIRQSPSRGLEWLGRTYFRSKWYN
DYAVSVKSRITINPDTSKNQFSLQLNSATPEDTAVYYCARQMDEGVGFDFWGQGTLVTVSS

$V_L$ EIVLTQSPATLSLSPGERATLSCRASQSVSSYLAWYQQKPGQAPRLLIYDASNRATGIPARFSG
SGSGTDFTLTISSLEPEDFAVYYCQQRSNWPPTFGQGTRLEIK

**ATA1.458.3**

$V_H$ QVQLQQSGPGLLKPSETLSLTCVISGDSVSTNDVTWNWIRQSPSRGLEWLGRTYFRRKWYN
DYARSVKSRITINPDTSKNQFSLQLNSATPEDTAVYYCARQMDEGVGFDFWGQGTLVTVSS

$V_L$ EIVLTQSPATLSLSPGERATLSCRASQSVSSYLAWYQQKPGQAPRLLIYDASNRATGIPARFSG
SGSGTDFTLTISSLEPEDFAVYYCQQRHNWPPTFGQGTRLEIK

**ATA1.458.5**

$V_H$ QVQLQQSGPGLLKPSETLSLTCVISGDSVSSNFVTWNWIRQSPSRGLEWLGRTYFRRKWYN
DYAVSVKSRITINPDTSKNQFSLQLNSATPEDTAVYYCARQMDEGVGFDFWGQGTLVTVSS

$V_L$ EIVLTQSPATLSLSPGERATLSCRASQSVSSYLAWYQQKPGQAPRLLIYDASNRATGIPARFSG
SGSGTDFTLTISSLEPEDFAVYYCQQRHSWPPTFGQGTRLEIK

**ATA1.459.3**

$V_H$ QVQLQQSGPGLLKPSETLSLTCVISGDSVSSNSVTWNWIRQSPSRGLEWLGRTYFRRRWYN
DYAVSVKSRITINPDTSKNQFSLQLNSATPEDTAVYYCARQMDKGVGFDFWGQGTLVTVSS

$V_L$ EIVLTQSPATLSLSPGERATLSCRASQSVSSYLAWYQQKPGQAPRLLIYDASNRATGIPARFSG
SGSGTDFTLTISSLEPEDFAVYYCQQRHNWPPTFGQGTRLEIK

**Tau-5**

$V_H$ EVQLQQSGAEIVRSGASVKLSCAASGFNIKDYYMHWVKQRPEQGLEWIGWIDPENGDIAY
APKFQGKATMTADTSSNTAYLQLSRLTSEDTAVYFCNGRGGMITTDFFDYWGQGTTLTVSS

$V_L$ DVLMTQTPLSLPVSLGDQASISCRSSQSIVHSNGNTYLEWYLQKPGQSPKLLIYKVSNRFSG
VPDRFSGSGSGTDFTLKISRVEAEDLGVYYCFQGSHVPWTFGGGTKLEIK

**zagotanemab**

$V_H$ EVQLVQSGAEVKKPGESLKISCKGSGYTFSNYWIEWVRQMPGKGLEWMGEILPGSDSIKYE
KNFKGQVTISADKSISTAYLQWSSLKASDTAMYYCARRGNYVDDWGQGTLVTVSS

$V_L$ EIVLTQSPGTLSLSPGERATLSCRSSQSLVHSNQNTYLHWYQQKPGQAPRLLIYKVDNRFSG
IPDRFSGSGSGTDFTLTISRLEPEDFAVYYCSQSTLVPLTFGGGTKVEIK

**Figure S1. Amino acid sequences of the tau antibody variable regions.** Soluble scFv-Fc fusion proteins (ATA1 series of antibodies) were expressed as VH-linker-VL-Fc with an IgG1 Fc. The amino sequence of the linker was GILGSGGGGSGG-GGSGGGGS. Tau-5 and zagotenemab were expressed as IgG1 proteins. For the mAbs, the first ten residues of the hinge and Fc region were EPKSCDKTHT, and for scFv-Fc fusions, the hinge and Fc region was truncated such that the first ten residues were KTHTCPPCPA. The last ten residues of the Fc region were QKSLSLSPGK for both mAbs and scFv-Fc fusions.

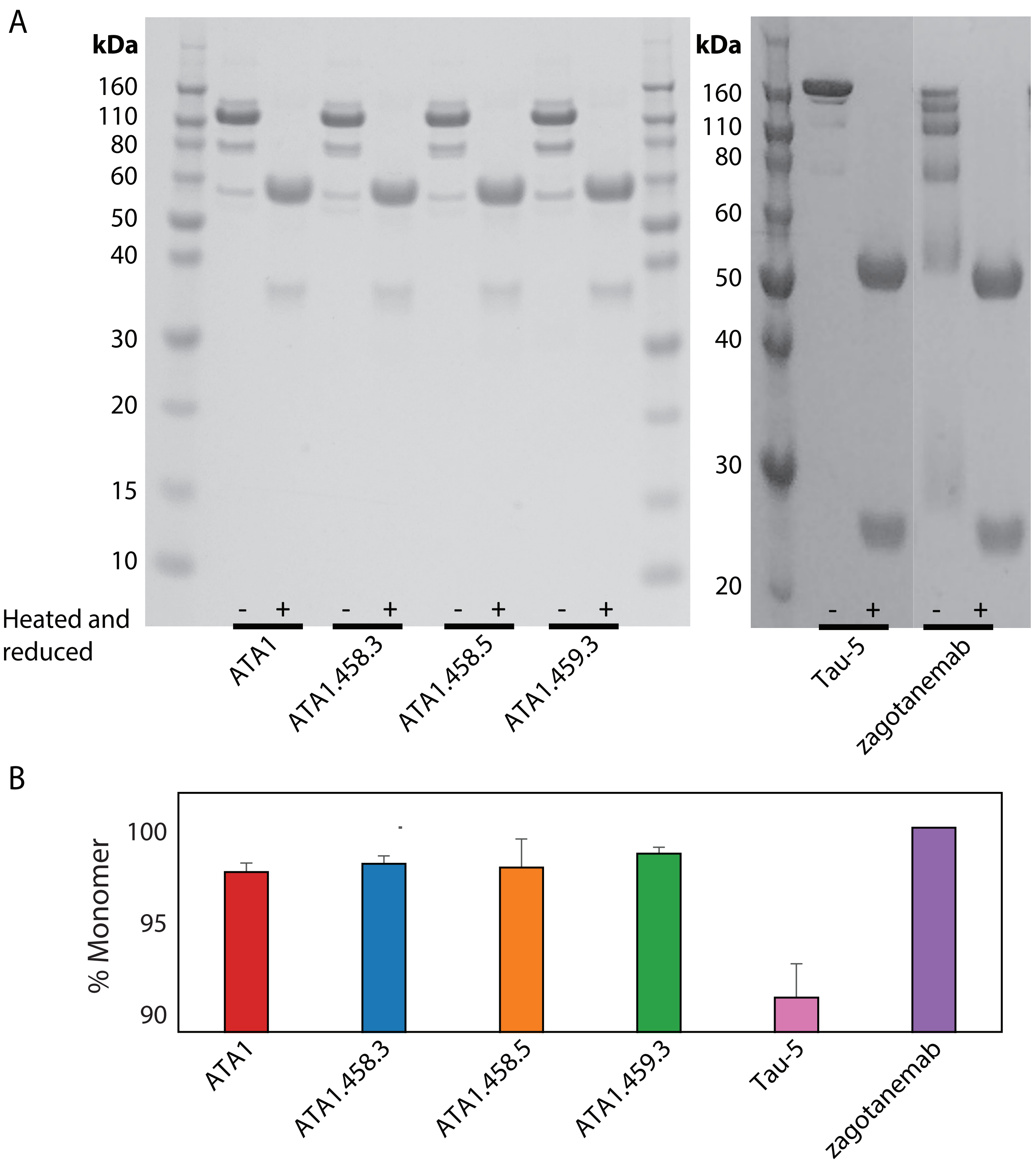


**Figure S2. Characterization of tau antibodies using SDS-PAGE and size-exclusion chromatography (SEC).** (A) SDS-PAGE analysis of antibody samples before (-) and after (+) being heated and reduced with β-mercaptoethanol. (B) SEC analysis of the percentage of antibody monomer following Protein A purification.

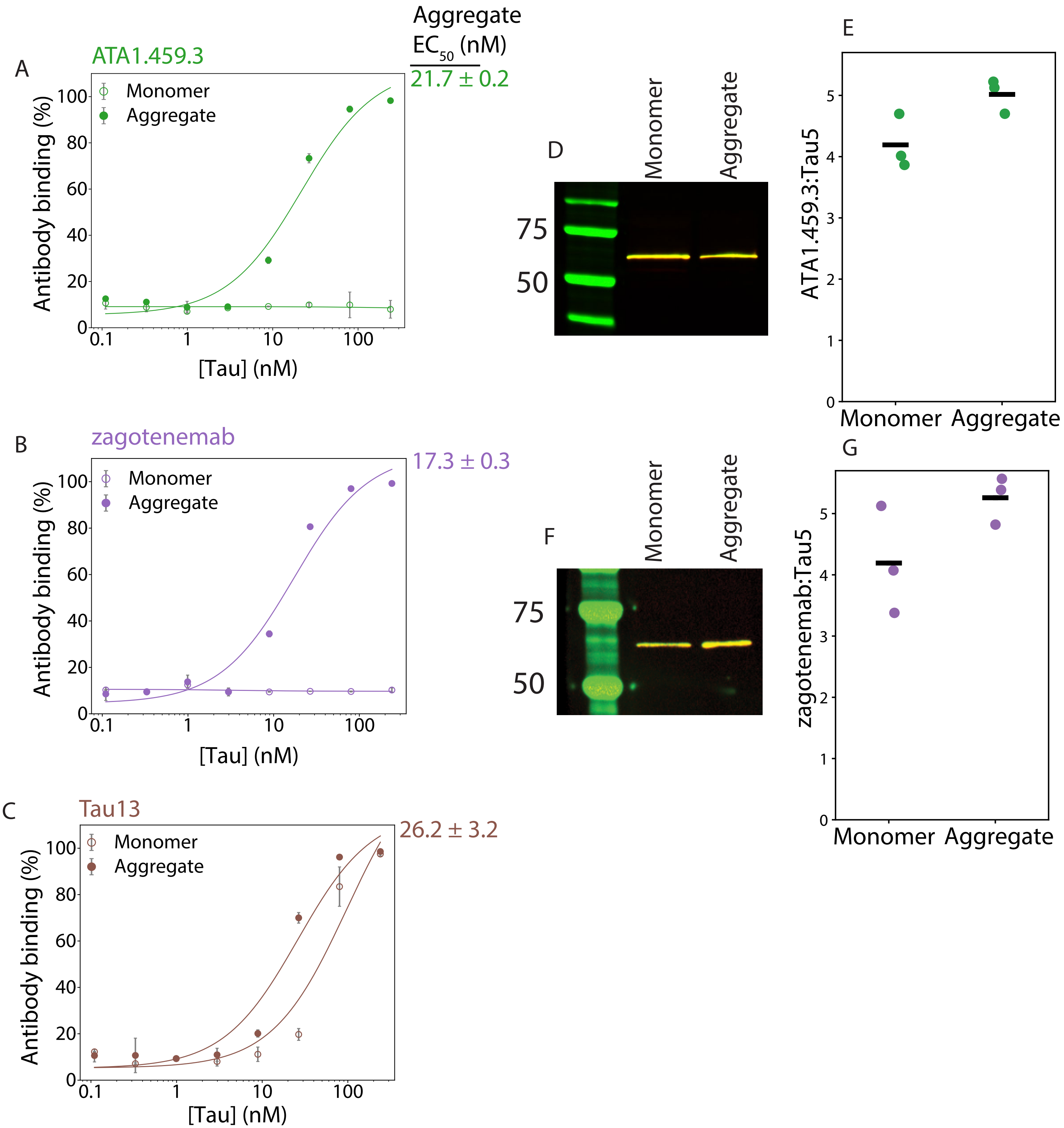


**Figure S3. Sandwich ELISA analysis of tau antibody conformational specificity.** (A) ATA1.459.3, (B) zagotenemab, and (C) Tau-13 were used as capture antibodies for recombinant tau monomer or tau fibrils. Captured tau monomer and fibrils were detected using the R1 polyclonal rabbit pan-tau primary antibody and a goat anti-rabbit HRP-conjugated secondary antibody. For (A-C), the data are the averages, and error bars are the standard deviations from three independent experiments. The binding of (E) ATA1.459.3 to recombinant tau monomer and fibrils was also examined via western blotting, and it was observed to bind tau monomer and fibrils under denaturing conditions. Binding was detected using goat anti-human Alexa Fluor 680 for ATA.459.3 and goat anti-mouse IRDye 800 for Tau-5. Three samples were analyzed, and a representative image is shown. (E) The data from three repeats were quantified for ATA1.459.3 binding to tau monomer and aggregates, and the average binding signals are shown as lines. (F) Zagotenemab showed similar binding to both tau monomer and tau aggregates via western blotting, and (G) quantification of this binding signal is shown for three repeats.

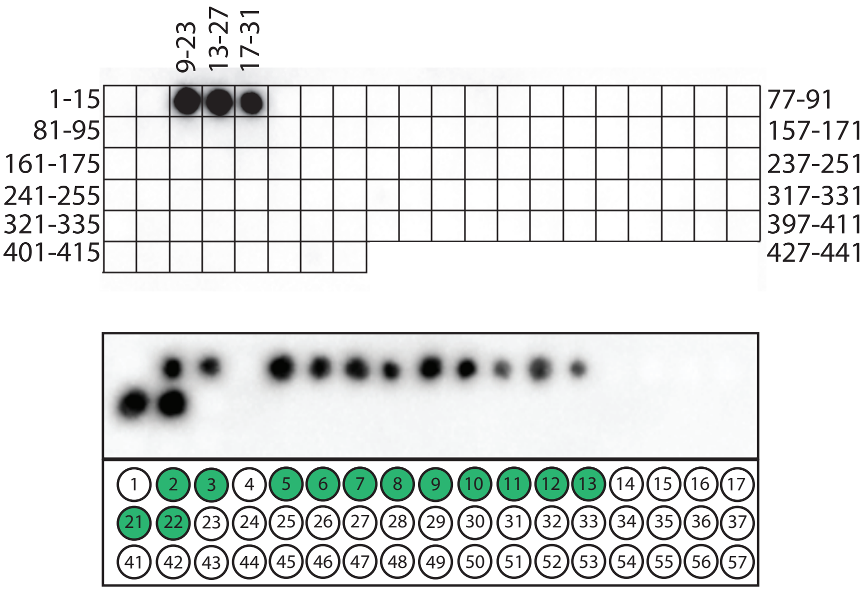


| | |
|---|---|
| 1 | RQEFEVMEDHAGTYG |
| 2 | QEFEVMEDHAG**TYGL** |
| 3 | EFEVMEDHAG**TYGL**G |
| 4 | FEVMEDHAGTYGLGD |
| 5 | EVMEDHAG**TYGL**GDR |
| 6 | VMEDHAG**TYGL**GDRK |
| 7 | MEDHAG**TYGL**GDRKD |
| 8 | EDHAG**TYGL**GDRKDQ |
| 9 | DHAG**TYGL**GDRKDQG |
| 10 | HAG**TYGL**GDRKDQGG |
| 11 | AG**TYGL**GDRKDQGGY |
| 12 | G**TYGL**GDRKDQGGYT |
| 13 | **TYGL**GDRKDQGGYTM |
| 14 | YGLGDRKDQGGYTMH |
| 15 | GLGDRKDQGGYTMHQ |
| 16 | LGDRKDQGGYTMHQD |
| 17 | GDRKDQGGYTMHQDQ |
| 21 | EVMEDHAG**TYGL**GDG |
| 22 | EVMEDHAG**TYGL**GGG |
| 23 | EVMEDHAGTYGGGGG |

**Figure S4. Peptide scanning analysis of ATA1.459.3 epitope.** (A) The epitope of ATA1.459.3 was first probed using a membrane containing 15-mer overlapping peptides for the full-length 2N4R tau protein, which are shifted in an interval of four amino acids. The epitope was identified as an N-terminal region using this first array. (B) A focused array containing overlapping tau 15-mer peptides from this N-terminal region, which differ by one amino acid, was then probed. This refined scanning identified a more focused epitope containing residues 16-TYGL-20 of the tau protein. Residues within the identified epitope are shown in green for peptides in which ATA1.459.3 binding was observed.

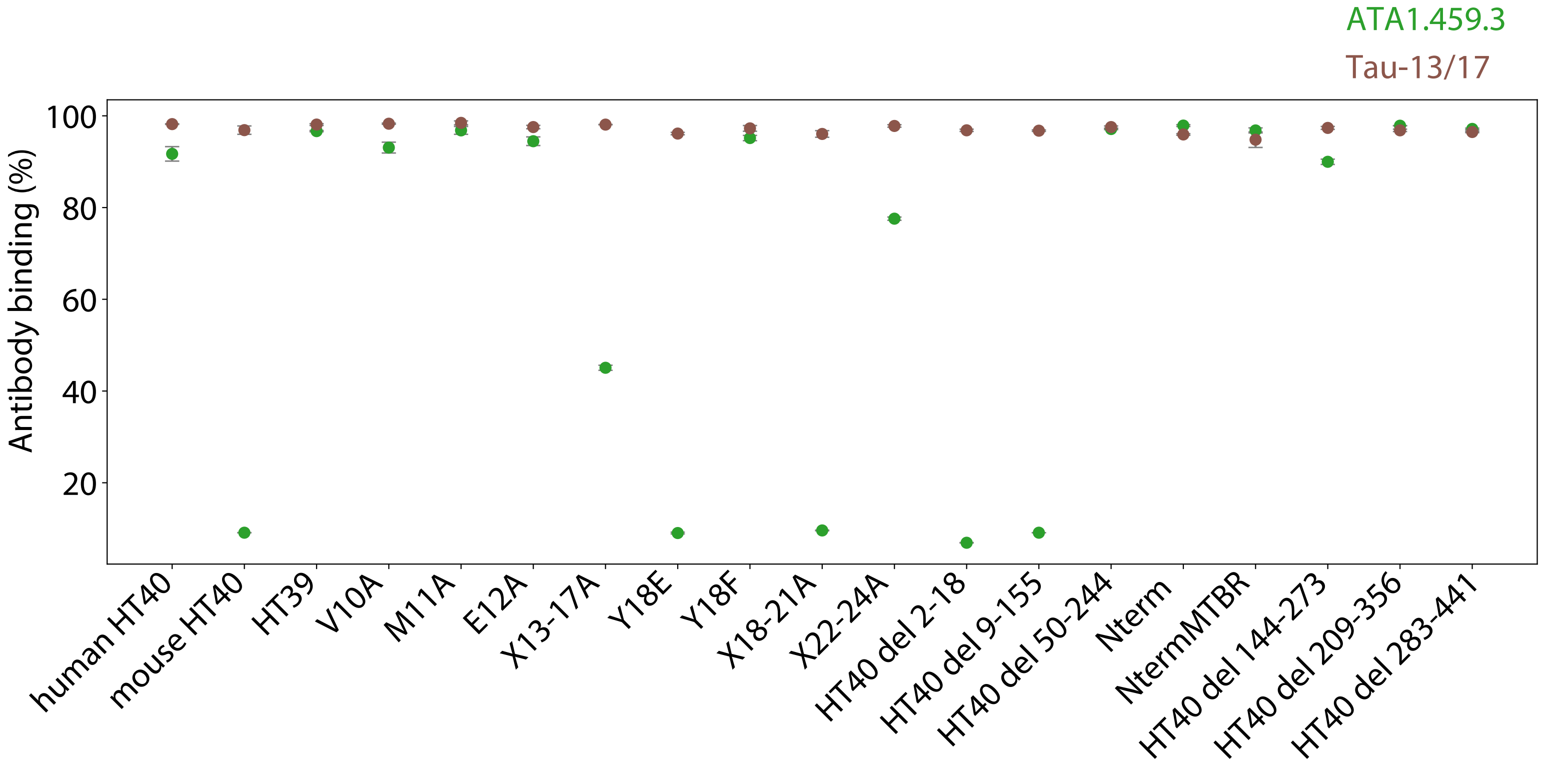


**Figure S5. ELISA analysis of ATA1.459.3 epitope.** The epitope of ATA1.459.3 was evaluated using protein fragments and protein mutants of the tau protein in an ELISA format. Tau protein fragments and mutants were immobilized in 96-well plates, and ATA1.459.3 or a mixture of Tau-13/Tau-17 antibody binding to immobilized tau proteins was detected with conjugated secondary antibody. Observed binding signals were converted to a linear scale as described in the Methods. Data are averages, and error bars are the standard deviations from two independent experiments.

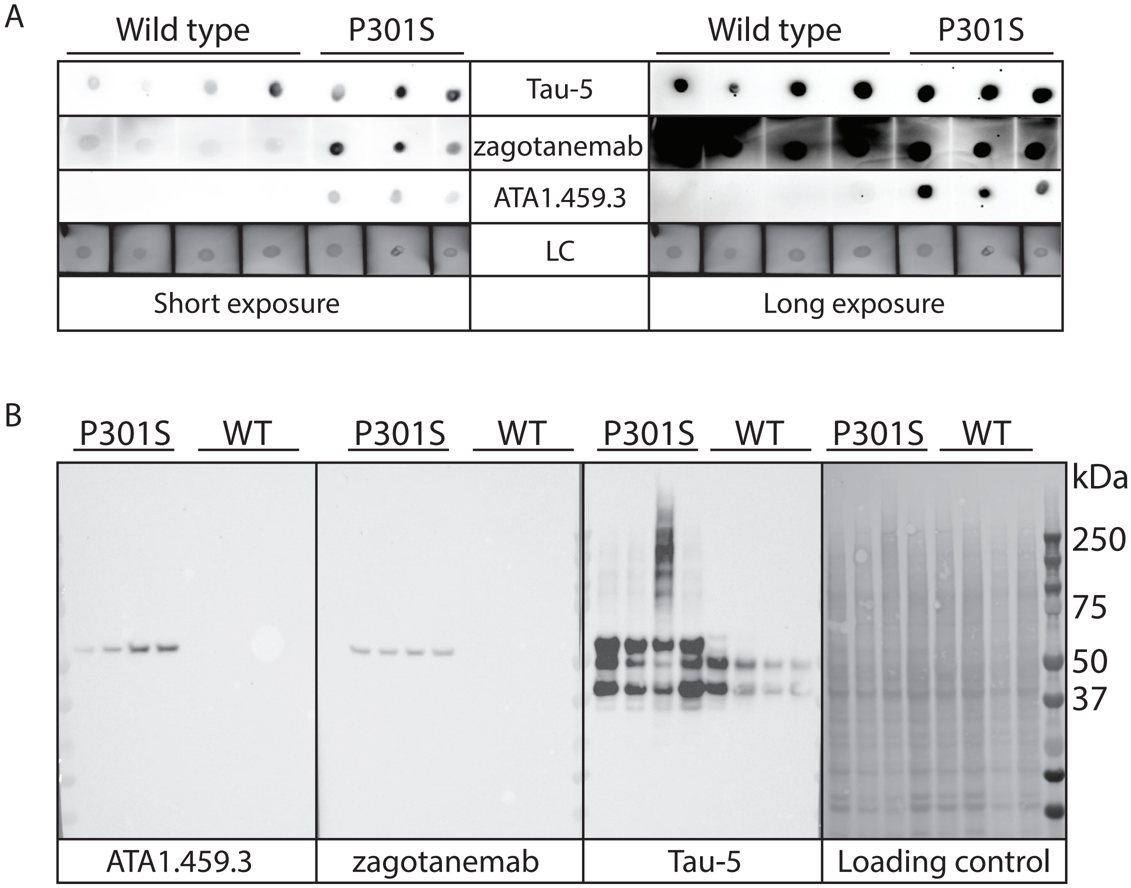


**Figure S6. Immunodot and western blot analysis of mouse brain lysates using tau conformational antibodies.** (A) Immunodot blotting analysis of wild-type and transgenic P301S tau mouse brains using ATA1.459.3, zagotenemab and Tau-5. Short (left) and long (right) time exposures are shown. The loading control (LC) was Ponceau stain. (B) Western blotting analysis of wild-type and P301S tau mouse brains using ATA1.459.3, zagotenemab, Tau-5, and Ponceau stain (loading control). In (A) and (B), the antibody binding was performed at 10 nM (1% milk in TBST overnight at 4 °C followed by anti-Fc HRP detection), the experiments were performed three times, and a representative example is shown.

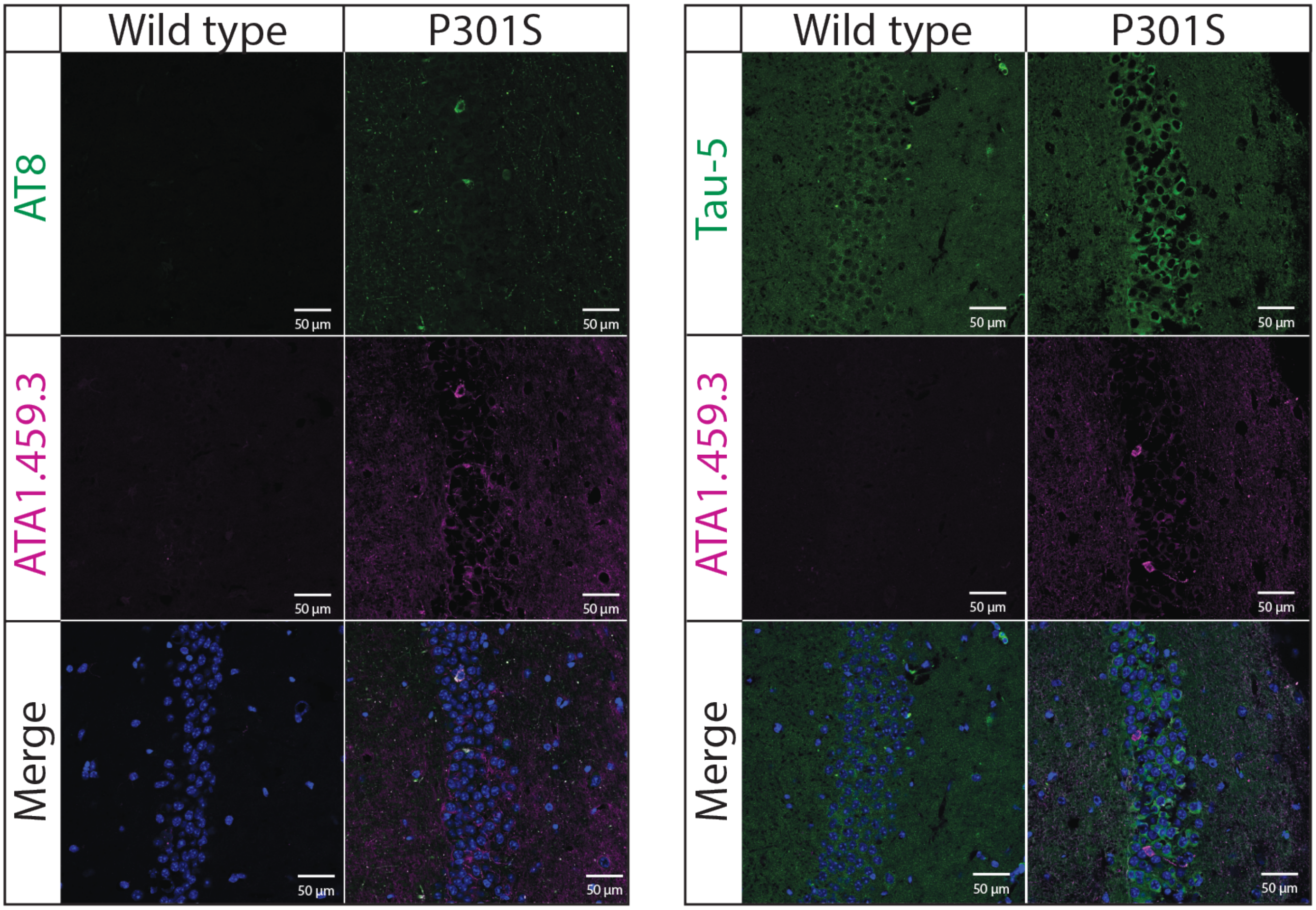


**Figure S7. Tau immunofluorescence analysis of brain tissues from transgenic P301S mice.** Fixed brain tissues from transgenic P301S or wild type mice were co-stained with a conformational tau antibody (ATA1.459.3, purple), a phospho-tau antibody (AT8, green) or a pan-tau antibody (Tau-5, green), and DAPI (blue). ATA1.459.3 was detected with anti-human Fc Alexa Fluor 647, and AT8 or Tau-5 were detected with anti-mouse Alexa Fluor 488. Individual fluorescence staining and merged results with DAPI are shown. The scale bars in the images are approximately 50 µm wide.

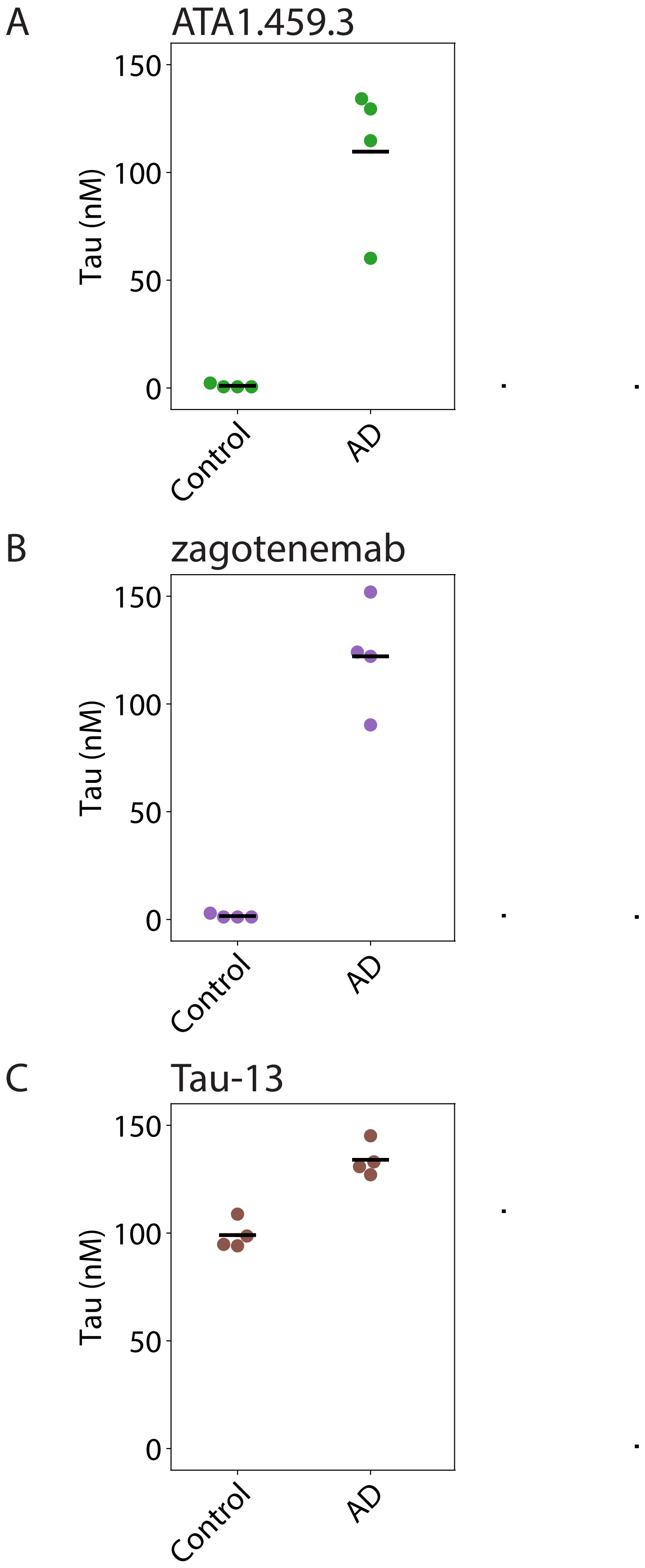


**Figure S8. Sandwich ELISA analysis of tau antibody binding to tau conformers in human brain lysates.** The binding of (A) ATA1.459.3, (B) zagotenemab, and (C) Tau-13 to human lysates from Alzheimer's disease (AD) and age-matched controls was examined via sandwich ELISA. ATA1.459.3, zagotenemab, and Tau-13 were immobilized in 96-well plates as capture antibodies, and lysate containing ~20 µg of total protein was added. Binding signal was detected using a polyclonal pan-tau primary rabbit antibody and goat anti-rabbit HRP-conjugated secondary antibody. The data are averages, and the error bars are standard deviations for two independent experiments.

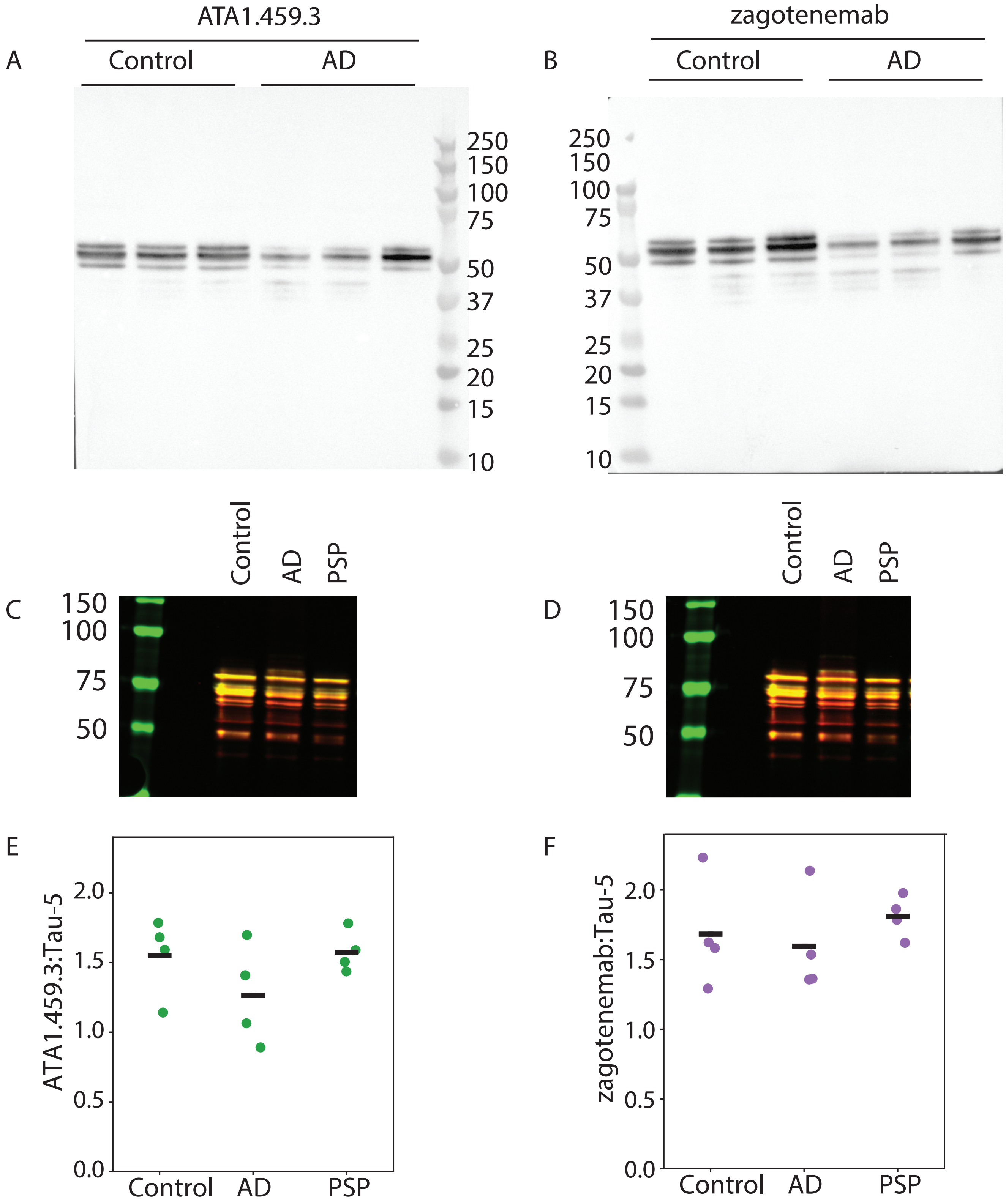


**Figure S9. Tau antibody western blotting analysis of human brain lysates.** The binding of (A) ATA1.459.3 and (B) zagotenemab to tau in human lysates from Alzheimer's disease (AD) and age-matched controls was examined via western blotting. Binding was detected using a goat anti-human HRP conjugated secondary antibody. Binding to all six isoforms of tau was observed for both ATA1.459.3 and zagotenemab in both AD and control tissues. The binding of (C) ATA1.459.3 and (D) zagotenemab to tau in human lysates from AD, progressive supranuclear palsy (PSP), and age-matched controls was examined in comparison to Tau-5 using fluorescent detection. Binding was detected using goat anti-human Alexa Fluor 680 for ATA.459.3 and zagotenemab, and goat anti-mouse IRDye 800 for Tau-5. The binding of both ATA1.459.3 and zagotenemab was observed to align with Tau-5 binding. Western blotting was performed using lysates obtained from 3-4 separate individuals for each diagnosis, and a representative image is shown. The binding signals observed for (E) ATA1.459.3 and (F) zagotenemab were quantified for each sample and is shown normalized to the signal for Tau-5. The lines represent the average signals.

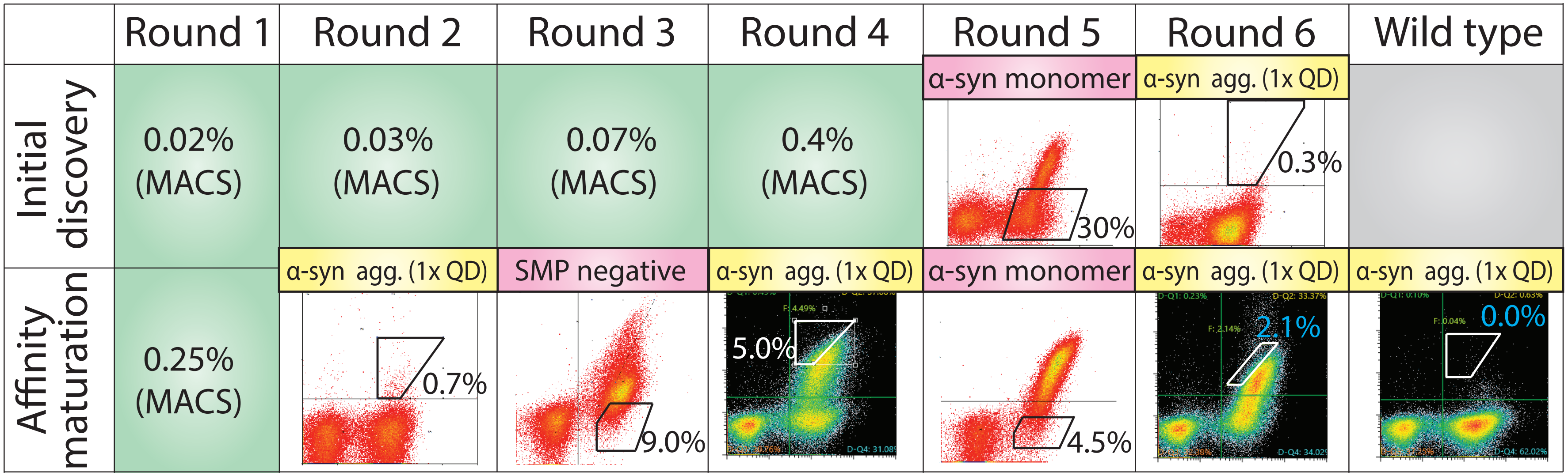


**Figure S10. Summary of library sorting to enrich for α-synuclein conformational antibodies.** In the initial discovery stage (top row), the human scFv library was enriched four times against α-synuclein fibrils via MACS (rounds 1-4; percentage of cells collected is reported), depleted once against α-synuclein monomer to remove non-conformational antibodies via FACS (round 5), and enriched one additional time against α-synuclein fibrils immobilized on QD immunoconjugates via FACS (round 6). A lead clone was then identified (aS2.1), and an scFab library was prepared via soft randomization of HCDR2. This library (bottom row) was then enriched against α-synuclein fibrils using MACS (round 1) and QD-fibril conjugates using FACS (round 2). A negative selection against a polyspecificity reagent (soluble membrane proteins) was then performed to deplete non-specific antibodies from the library (round 3). Next, a positive selection was performed by FACS using QD-fibril conjugates (round 4), a negative selection was performed by FACS against α-synuclein monomer (round 5), and a final positive selection was performed by FACS using QD-fibril conjugates. The terms for α-synuclein in the figure refer to fibrils (α-syn agg.) and α-synuclein monomer. The percentages in the green boxes represent the percentage of cells collected in MACS selections, and the percentages in the cytogram gates represent the percentage of cells collected in the FACS selections. Blue numbers represent the % of cells collected for the library and wild-type samples that were evaluated at the same time, which can be directly compared.

**aS2.1 WT**

$V_H$ QVQLQQSGPGLVKPSQTLSLTCAISGDSVSSHSASWNWIRQSPSRGLEWLGRTYYRSKWYN
DYAVSVKSRIIINPDTSKNQFSLQLDSVTPEDTAVYFCTRATRGASDYWGQGTLVTVSS

$V_L$ EIVLTQSPGTLSLSPGERATLSCRASQSVSSSYLAWYQQKPGQAPRLLIYGASNRATGIPDRFS
GSGSGTDFTLTISRLEPEDFAVYYCQQYGSSPQTFGQGTKVEIK

**H2.3**

$V_H$ QVQLQQSGPGLVKPSQTLSLTCAISGDSVSSHSASWNWIRQSPSRGLEWLGRTYYRGKWKT
DYAVSVKSRIIINPDTSKNQFSLQLDSVTPEDTAVYFCTRATRGASDYWGQGTLVTVSS

$V_L$ EIVLTQSPGTLSLSPGERATLSCRASQSVSSSYLAWYQQKPGQAPRLLIYGASNRATGIPDRFS
GSGSGTDFTLTISRLEPEDFAVYYCQQYGSSPQTFGQGTKVEIK

**H2.4**

$V_H$ QVQLQQSGPGLVKPSQTLSLTCAISGDSVSSHSASWNWIRQSPSRGLEWLGRTYYRSKWHN
DYAASVKGRIIINPDTSKNQFSLQLDSVTPEDTAVYFCTRATRGASDYWGQGTLVTVSS

$V_L$ EIVLTQSPGTLSLSPGERATLSCRASQSVSSSYLAWYQQKPGQAPRLLIYGASNRATGIPDRFS
GSGSGTDFTLTISRLEPEDFAVYYCQQYGSSPQTFGQGTKVEIK

**H2.7**

$V_H$ QVQLQQSGPGLVKPSQTLSLTCAISGDSVSSHSASWNWIRQSPSRGLEWLGRTYYRGKWRV
DYAFSVKSRIIINPDTSKNQFSLQLDSVTPEDTAVYFCTRATRGASDYWGQGTLVTVSS

$V_L$ EIVLTQSPGTLSLSPGERATLSCRASQSVSSSYLAWYQQKPGQAPRLLIYGASNRATGIPDRFS
GSGSGTDFTLTISRLEPEDFAVYYCQQYGSSPQTFGQGTKVEIK

**cinpanemab**

$V_H$ EVQLVESGGGLVEPGGSLRLSCAVSGFDFEKAWMSWVRQAPGQGLQWVARIKSTADGGTT
SYAAPVEGRFIISRDDSRNMLYLQMNSLKTEDTAVYYCTSAHWGQGTLVTVSS

$V_L$ SYELTQPPSVSVSPGQTARITCSGEALPMQFAHWYQQRPGKAPVIVVYKDSERPSGVPERFSG
SSSGTTATLTITGVQAEDEADYYCQSPDSTNTYEVFGGGTKLTVL

**Figure S11. Amino acid sequences of the α-synuclein antibody variable regions.** The antibodies were expressed as IgG1s, as defined in Fig. S1.

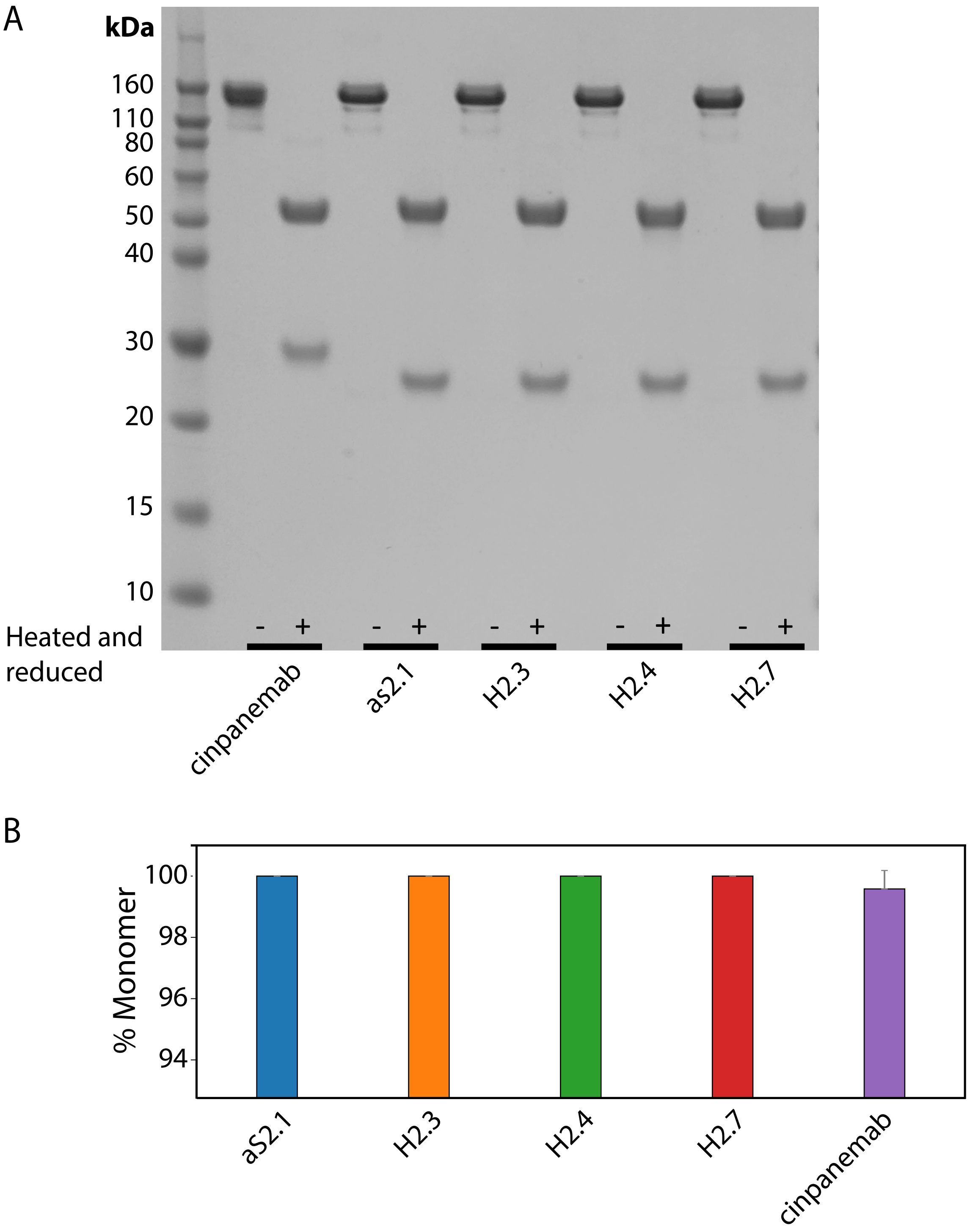


**Figure S12. Characterization of α-synuclein antibodies using SDS-PAGE and SEC.** (A) SDS-PAGE analysis of antibodies before (-) and after (+) being heated and reduced with β-mercaptoethanol. (B) SEC analysis of the percentage of antibody monomer following Protein A purification.

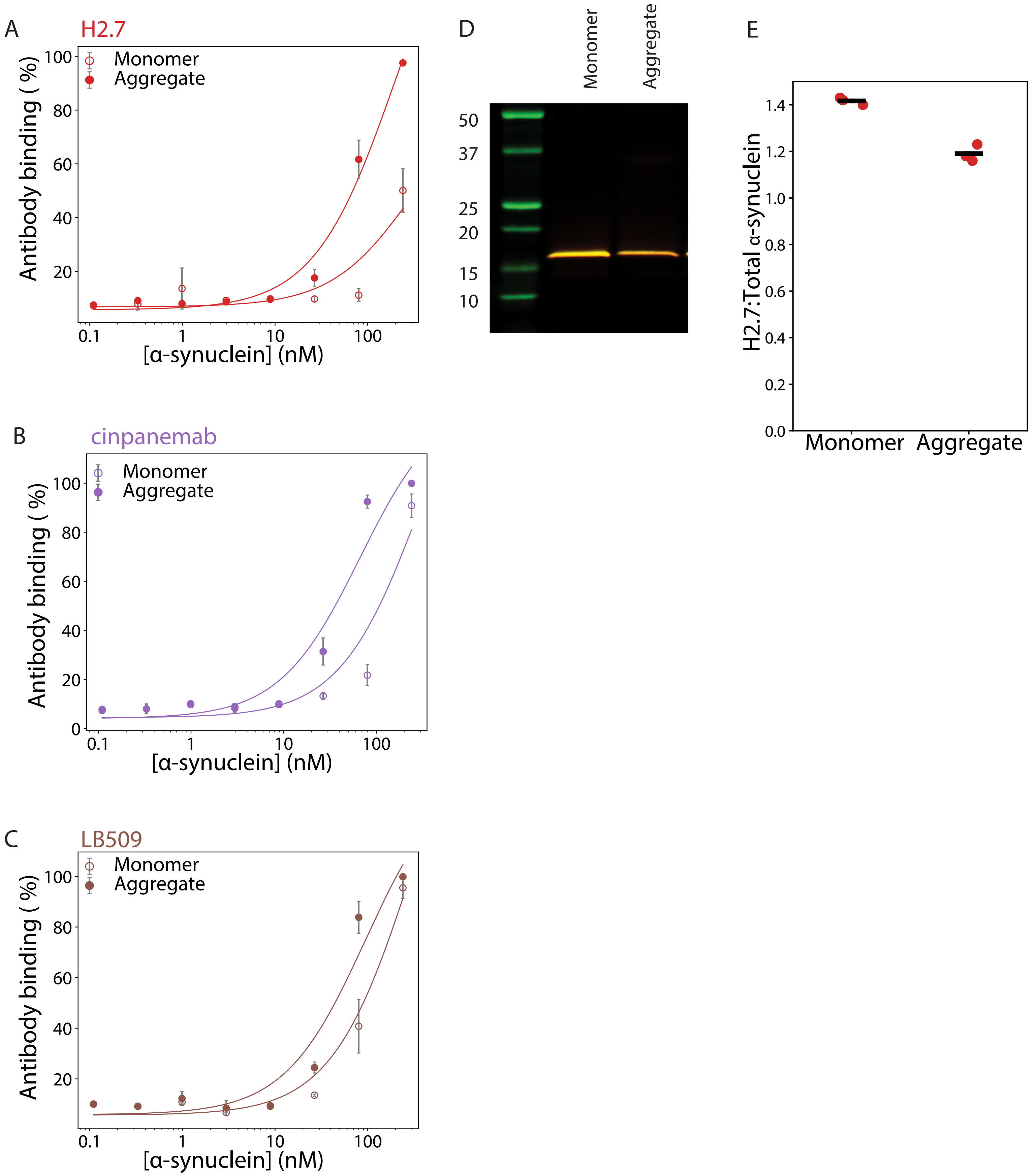


**Figure S13. Sandwich ELISA analysis of α-synuclein antibody conformational specificity.** (A) H2.7, (B) cinpanemab, and (C) LB509 were used as capture antibodies for recombinant α-synuclein monomer or fibrils. Captured monomer and fibrils were detected using a monoclonal rabbit anti-α-synuclein primary antibody and a goat anti-rabbit HRP-conjugated secondary antibody. For (A-C), the data are averages, and error bars are the standard deviations for three independent experiments. The binding of (D) H2.7 to recombinant α-synuclein monomer and fibrils was also examined with western blotting, and it was observed to bind α-synuclein monomer and fibrils under denaturing conditions. Three repeats were performed, and a representative image is shown. (E) The data was quantified compared to a total α-synuclein antibody. The lines represent the average binding signal.

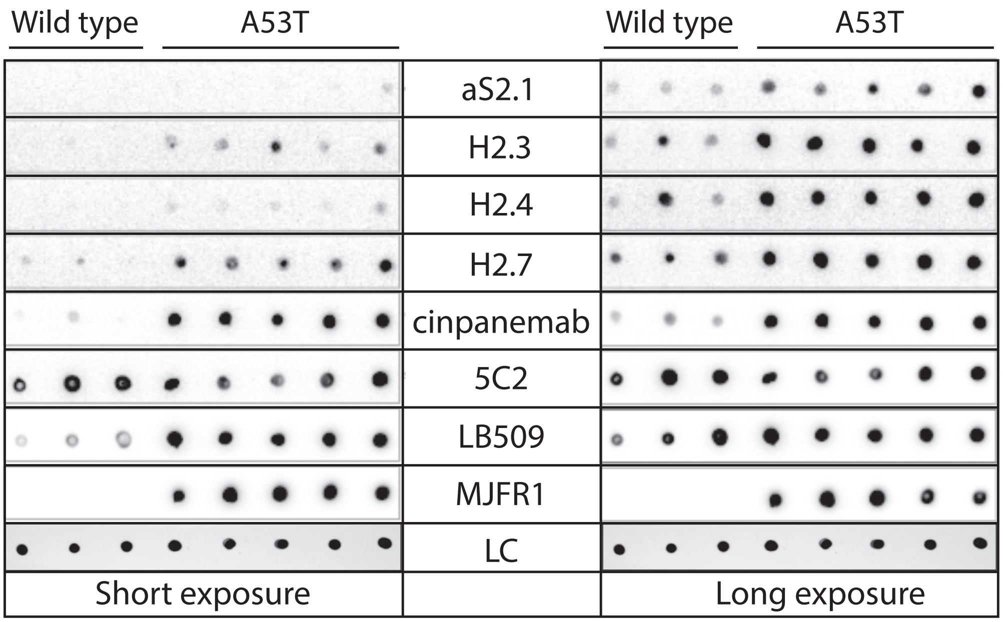


**Figure S14. Immunodot blot analysis of brain samples from wild-type and α-synuclein transgenic (A53T) mice.** (A) Immunodot blotting analysis of wild-type and transgenic (A53T) α-synuclein mouse brains. Short (left) and long (right) time exposures are shown. The loading control (LC) was Ponceau stain, and the antibody binding conditions were the same as those in Fig. S6A. The experiments were performed three times and a representative example is shown.

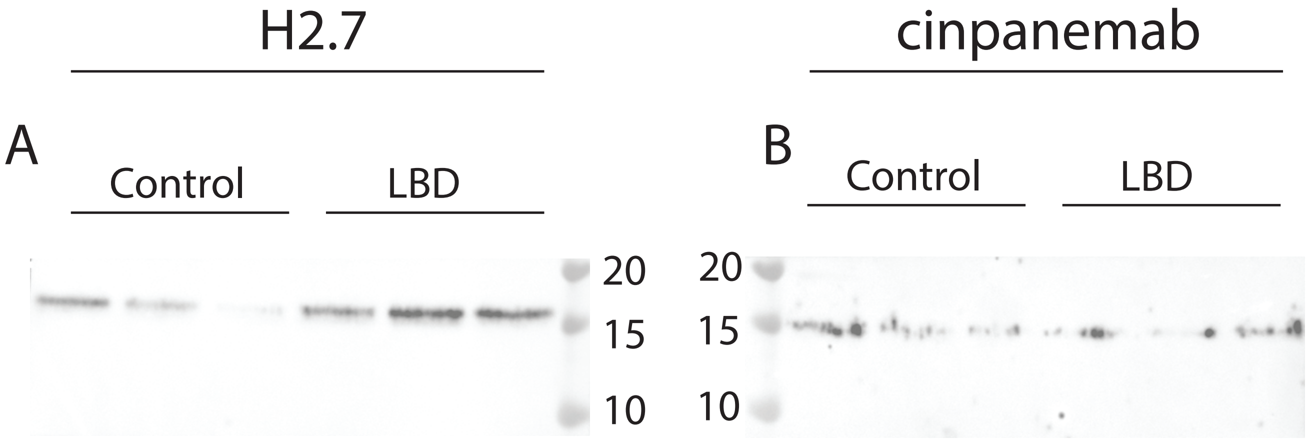


**Figure S15. α-synuclein antibody western blotting analysis of human brain lysates.** The binding of (A) H2.7 and (B) cinpanemab to α-synuclein in human lysates from Lewy Body Dementia (LBD) and age-matched controls was examined via western blotting. Binding was detected using a goat anti-human HRP conjugated secondary antibody. Binding to α-synuclein was observed for both H2.7 and cinpanemab in both LBD and control tissues. The binding of (C) H2.7 to recombinant α-synuclein monomer and fibrils was also examined with western blotting, and it was observed to bind α-synuclein monomer and fibrils under denaturing conditions.

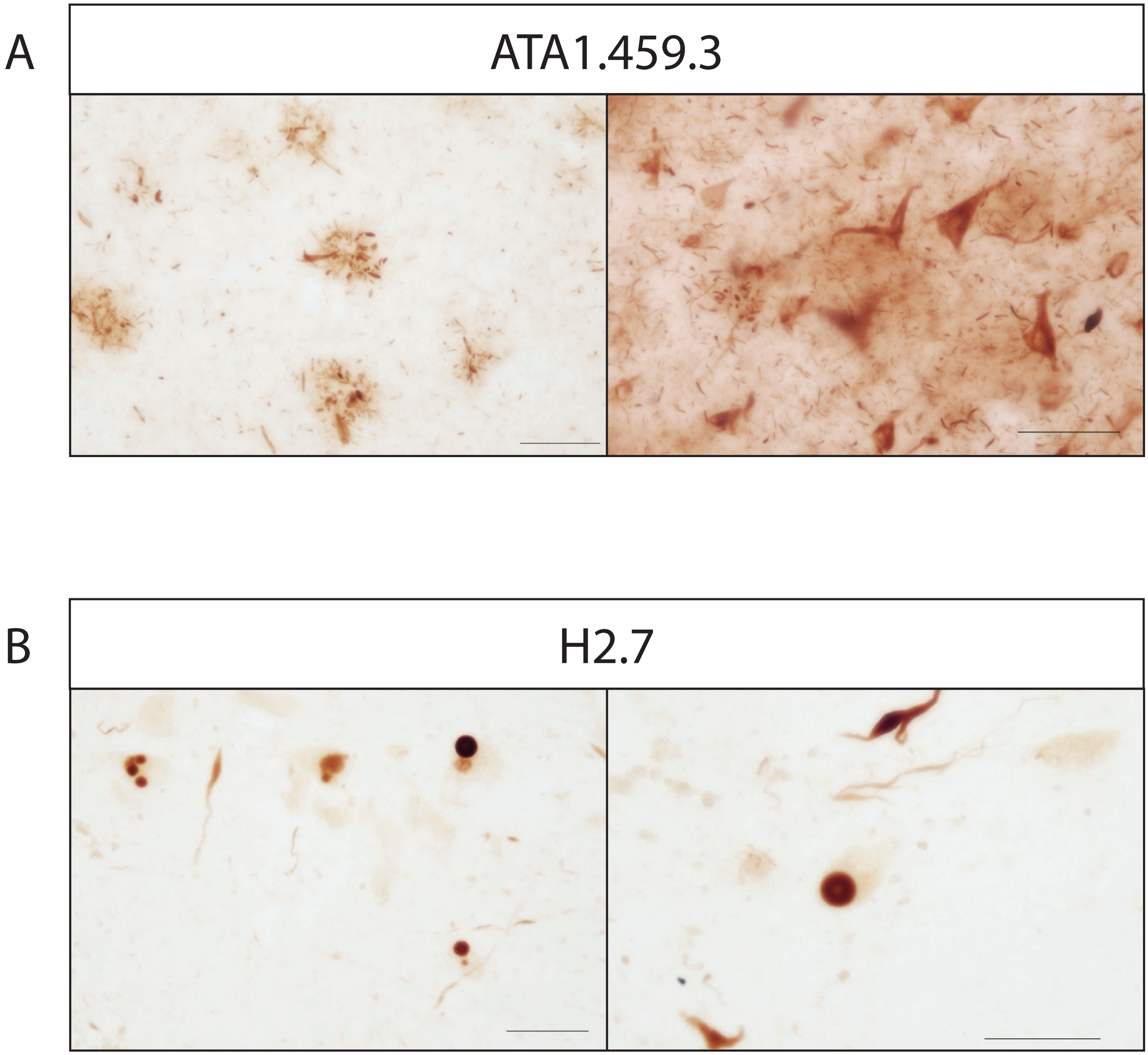


**Figure S16. Immunohistochemical staining of free-floating human brain tissue.** Antibody staining to free-floating human tissue samples was examined in the absence of antigen retrieval. (A) ATA1.459.3 binding to samples from the inferior temporal gyrus of Alzheimer's disease cases was examined. Binding of ATA1.459.3 to aggregated tau was observed in these samples. (B) H2.7 binding to samples from the substantia nigra of Parkinson's disease cases was examined. Binding of H2.7 to α-synuclein aggregates, including Lewy bodies, was observed in these samples.

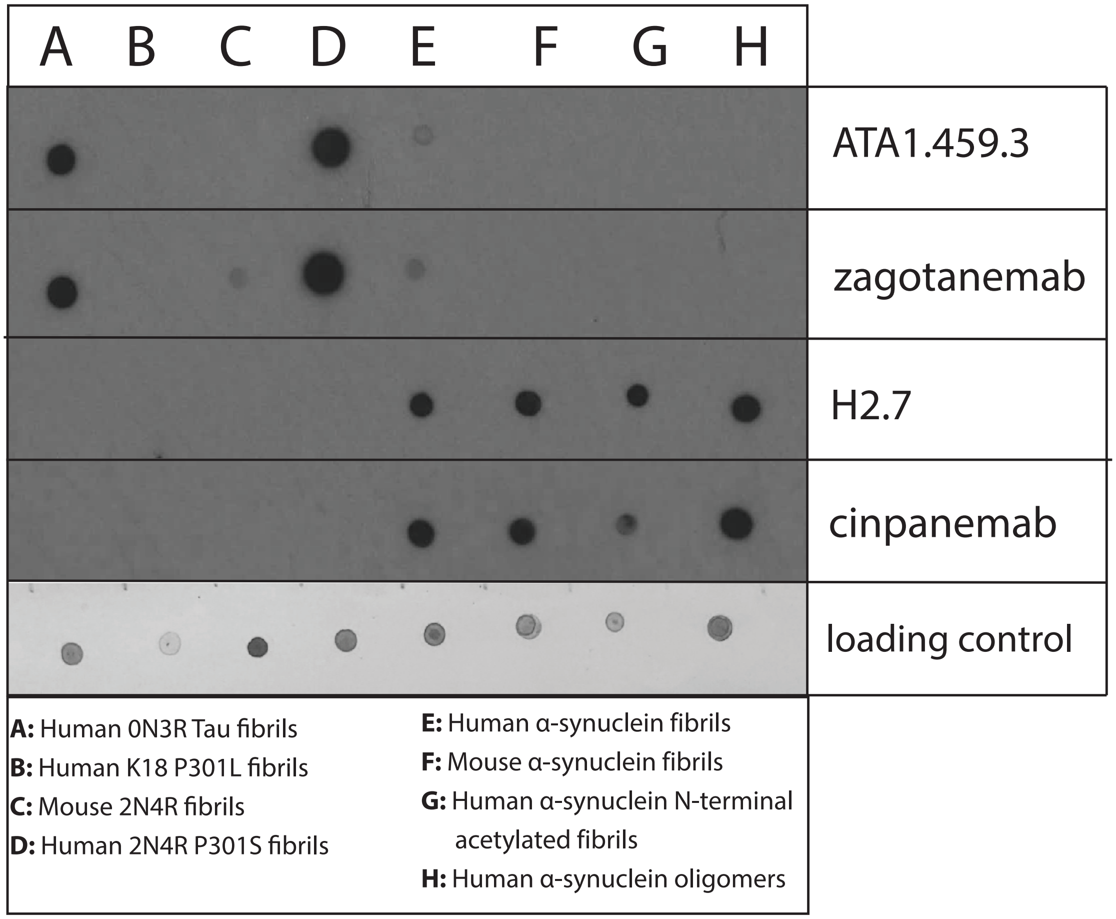


**Figure S17. Immunoblotting analysis of tau and α-synuclein antibody sequence specificity.** The binding of ATA1.459.3, zagotenemab, H2.7, and cinpanemab to various conformers of tau and α-synuclein was examined via immunoblotting. Binding of ATA1.459.3 and zagotenemab was observed to human 2N4R tau fibrils as well as human 0N3R fibrils. Binding of H2.7 and cinpanemab was observed to human α-synuclein oligomers as well as human and mouse α-synuclein fibrils. Loading was detected via silver stain. Immunoblotting was repeated twice, and a representative image is shown.

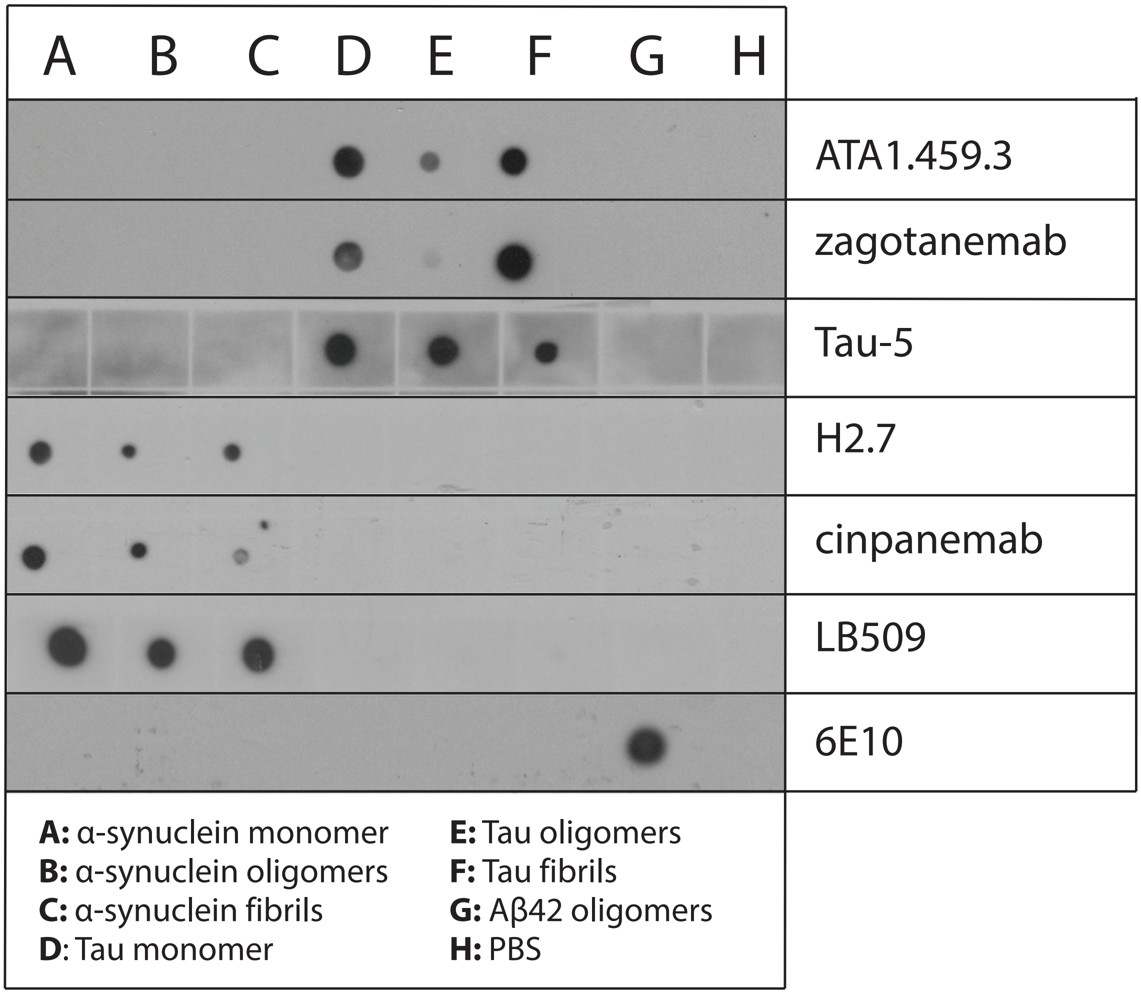


**Figure S18. Immunoblotting analysis of tau and α-synuclein antibody specificity.** The binding of ATA1.459.3, zagotenemab, H2.7, and cinpanemab to tau and α-synuclein monomer, oligomers, and fibrils and Aβ oligomers was examined via immunoblotting. Binding of ATA1.459.3 was observed for tau monomer, oligomers, and fibrils, and binding of zagotenemab was observed for both tau monomer and fibrils. Tau loading was confirmed with Tau-5 staining. Binding of H2.7 and cinpanemab was observed for α-synuclein monomer, oligomers, and fibrils. α-synuclein loading was confirmed with LB509 staining. Aβ oligomer loading was confirmed with 6E10 staining.
